# Blending physics-based and inverse folding models to disentangle variant effects on stability and function

**DOI:** 10.64898/2026.07.30.741764

**Authors:** Ezequiel A. Galpern, Xavier Soler Sanchis, Charles W. J. Pugh, Federico Billeci, Jonathan Frazer, Mafalda Dias

**Affiliations:** Centre for Genomic Regulation (CRG), The Barcelona Institute of Science and Technology, Dr. Aiguader 88, Barcelona 08003, Spain; University Pompeu Fabra (UPF), Barcelona, Spain

## Abstract

Protein sequences are constrained not only by the need to fold into stable structures, but also by specific functional requirements imposed by natural selection. Yet predictions of how amino-acid changes affect proteins typically collapse these constraints into a single scalar score. Quantitatively separating these effects at scale remains an open challenge, with direct relevance spanning protein design to understanding the molecular mechanisms of disease. Inverse-folding (IF) models have emerged as fast, unsupervised predictors of folding energy changes (ΔΔG), but because they learn statistical correspondences between structure and sequence, they can conflate conservation driven by function with conservation driven by stability. Here, we show that blending IF models with a physics-based coarse-grained potential improves global correlation with experimental ΔΔG and, crucially, reduces IF model bias at functional sites. Applying the best-performing blend together with an evolutionary language model, we decompose each variant’s evolutionary cost into folding energy and dark energy, the latter capturing functional constraints beyond folding stability. With this decomposition, and without the need for supervision, we find that disease gain-of-function variants show a distinct functional signature from loss-of-function variants. In particular, we identify oncogenic drivers as largely preserving stability while exhibiting high dark energy, as opposed to tumor suppressors which are predominantly destabilized, paving the way to a mechanistic understanding of driver mutations in cancer. Together, these results provide a scalable framework for accurate ΔΔG prediction and mechanistic disentanglement of variant effects.

## Introduction

Natural selection acts on the many functions a protein must perform, from binding specific ligands or catalysing reactions, and encodes the information needed to sustain each of them in the genome. Folding stability is one such constraint: it has shaped protein sequences throughout evolutionary history and distinguishes them from random heteropolymers [1,2]. But while folding is often necessary, it is not sufficient for function. The precise chemistry required for catalysis or binding is also imprinted in sequences, and this chemistry can be neutral or even detrimental to stability [3], revealing a trade-off between stability and function [4]. Disentangling these distinct constraints is essential both for designing proteins with improved or novel functions and for understanding the molecular mechanisms underlying biological phenomena, including in disease and health.

High-throughput assays such as deep mutational scanning (DMS) attempt to systematically quantify the effects of sequence variation, but the measured phenotypes are often a proxy for molecular properties, such as chemical activity, protein abundance, or organismal fitness [5]. Most of these quantities fold several, sometimes competing, evolutionary constraints into a one-dimensional score, and so do not by themselves identify the protein-level mechanism through which a variant acts [6]. The situation is even more extreme in our current understanding of the clinical impact of human missense variants, which are often reduced to a summary label, whether pathogenic versus benign, or loss-of-function versus gain-of-function [7]. Multiplexed-readout DMS can begin to separate different mechanistic contributions, but this approach remains experimentally intractable at proteome scale [8–13].

Because folding is a major evolutionary constraint, evolutionary and folding energy landscapes agree globally, but they can disagree locally. The difference between the evolutionary free energy and the physical folding free energy has been defined as “dark energy,” which quantifies additional evolutionary constraints beyond folding [14]. Local perturbations to the dark energy can be computed variant by variant, by subtracting the folding free-energy change (experimentally measured as ΔΔ*G*) from the evolutionary free energy change predicted by a protein language model. Other approaches have been proposed to isolate functional contributions from evolutionary scores at the proteome scale [15,16]. In practice, however, all these decompositions rely on high-throughput ΔΔ*G* data or, when unavailable, on accurate and unbiased computational predictions of folding-stability changes.

In recent years, inverse-folding (IF) models have emerged as a scalable route to predicting ΔΔ*G* [17,18]. IF models learn amino-acid sequence propensities conditioned on a given backbone, and can therefore generate de novo sequences compatible with a target structure. Models such as ESM-IF1 and ProteinMPNN have also been used as unsupervised zero-shot predictors of ΔΔ*G*, although the formal connection between thermodynamic stability and inverse-folding probabilities is still being studied [19]. More recently, correction schemes have been proposed to improve IF models’ stability prediction [20], alongside models such as PottsMPNN that move beyond native sequence recovery toward improved modeling of sequence-energy landscapes [21]. Training on ensembles of structures has likewise been introduced to average-out non-structural evolutionary signals, such as phylogenetic relatedness, that a model might otherwise memorize [22].

However, sequence compatibility with a structure is not necessarily a pure folding-stability signal: functional residues can be conserved because of catalysis, binding, allostery or regulation [23,24], and all these evolutionary constraints coexist in natural sequences. Even if the natural structure-sequence mapping were learned perfectly, zero-shot IF ΔΔ*G* predictions could still be contaminated by projections of non-folding functional signals [25]. Physics-based ΔΔ*G* predictors offer a complementary route, estimating mutation-induced stability changes from protein structures using explicit energetic models. In particular, AWSEM is a coarse-grained, transferable potential grounded in energy landscape theory and previously used in frustration and dark-energy analyses of protein structure, sequence and function [14,26–28]. Here, we use AWSEM as a physical counterpart to IF models, testing whether the two capture complementary information about mutational stability effects, particularly in functionally relevant sites. We then use the resulting folding-stability estimates to disentangle the molecular effect of disease-associated variants into folding and dark-energy components.

## Results

### Blending Inverse Folding and physics-based models improves global correlation with ΔΔG experimental data

We developed a strategy to approximate the folding free-energy change ΔΔ*G* caused by single-site mutations. We leveraged existing models of two different kinds, structure-conditioned protein sequence models that capture the structure-to-sequence statistical correspondence and physics-based approaches. To exploit their complementarity, we defined a blend of their mutation-level energy change predictions for the same protein structure, sequence, and variant.

On the one hand, we considered a force-field rooted in the energy landscape theory, the Associative Memory Water Mediated, Structure and Energy Model (AWSEM) [26]. This physics-based coarse-grained, transferable potential has been extensively used in frustration analysis for localizing protein regions required for folding and function [24,27], and recently for measuring the folding and dark energy changes for sequence variants [14]. Given a structure *S* and a sequence *s*, the AWSEM potential delivers an estimation of the energy gap *E*(*S, s*) between the native folded state and a molten-globule reference state. Neglecting the entropic change, upon a single site variant we take Δ Δ *G* (*S, s* → *s*′) Δ *E* (*S,s* → *s*′).

On the other hand, we used the Inverse Folding (IF) models ESM-IF1 [17], ProteinMPNN [18] and PottsMPNN [21]. ESM-IF1 and ProteinMPNN estimate a probability distribution over sequences conditioned on a structure *S*. The effective energetic penalty for changing an amino acid in a given backbone Δ *E* (*S,s* → *s*′) = − [*log P* (*s*′ | *S*) − *log P* (*s* | *S*)] has been used as a proxy for ΔΔ*G* showing a strong empirical correlation with experimental values [5,17]. PottsMPNN is a recently developed method that converts ProteinMPNN encoding into a structure-specific Potts model and incorporates a training step using multiple sequence alignments [21]. The corresponding mutation score Δ *E* (*S, s* → *s* ′) can therefore be obtained by evaluating the Potts model directly.

We computed the folding energy change Δ *E*^*fold*^ (*S, s* → *s*′) as a blend of two different models, while reserving ΔΔ*G* strictly for experimental values. We defined a blend as a scale-matched weighted sum of the scores obtained with AWSEM and an IF model (Fig. 1A),

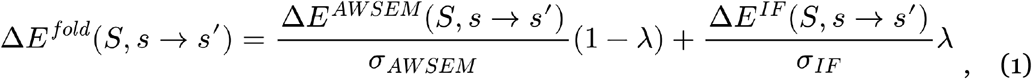

where λ is the blend weight and σ is the average standard deviation of a Δ*E* model over all possible single-site variants of a reference protein set that sets the arbitrary units of that model to a common scale (see *Methods*). This scale matching is motivated by empirical observations that the standard deviation of ΔΔ*G* is nearly constant across protein families [29,30] and by extending the argument to the model scores. Also, following the same definition, Δ *E*^*fold*^ can be rescaled to kcal/mol by multiplying it to λ _ΔΔ*G*_ if available.

**Figure 1.**
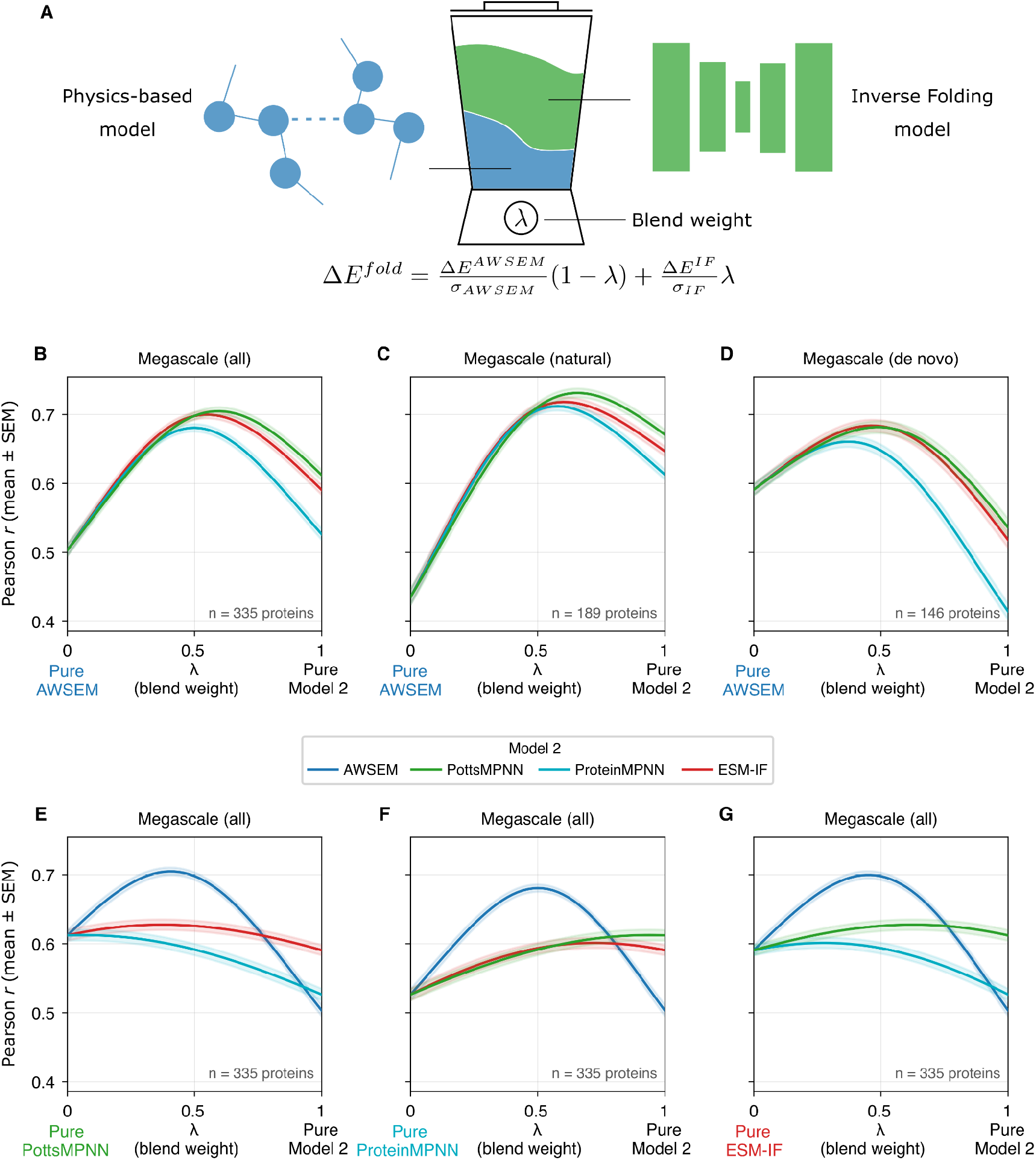
Blend model definition and global performance. **A**. AWSEM-IF blends estimate the folding stability gap change induced by single-site variants by a scale-matched weighted average of the predictions of a physics-based model, AWSEM and an Inverse Folding (IF) model. **B**. Correlation between AWSEM-IF blend model predictions and the Megascale experimental data measured with the Pearson’s r coefficient mean over all single-site variants of 335 natural and de novo proteins and its standard error (SEM), as a function of the blend weight, for AWSEM-PottsMPNN (green), AWSEM-ProteinMPNN (purple) and AWSEM-ESM-IF (red). **C-D**. Correlations for only the 189 natural proteins (C) and for only the de novo designs (D). **E-G**. Correlations for all possible blend combinations of the IF and AWSEM models. IF-IF blends underperformed AWSEM-IF blends.

We compared Δ *E*^*fold*^ to ΔΔ*G* experimental values for 335 proteins of the Rocklin’s Megascale dataset [31]. We considered single-site variants for both the natural proteins and the de novo designs of the dataset and computed the correlation between Δ *E*^*fold*^ and ΔΔ*G* for each protein. Although AWSEM alone underperforms all IF models, blending with AWSEM significantly improves every IF model, with the highest correlations at intermediate blend weights (Fig. 1B, Table S1). The improvement remains significant in both natural proteins and de novo designs (Fig. 1C,D), though pure IF models perform worse than pure AWSEM on de novo designs. When we generalize the strategy to blend between pairs of IF models, the improvement is marginal compared to blends that include the physics-based force field AWSEM (Fig. 1E-G), highlighting the fact that performance improvement comes from complementary information between structure-conditioned protein-sequence and physics-based models. We repeated the analysis using single-residue corrections from Dutton et al. for ProteinMPNN and ESM-IF1, which improves the performance of the pure models for ΔΔ*G* prediction [20], and find that the blends with AWSEM also benefit from them (Fig. S1).

### Blended models recover folding stability change prediction at functional sites

We next asked whether the improved global ΔΔ*G* prediction obtained by blending AWSEM with IF models reflects a correction of systematic local errors, particularly at functional sites. Many protein functional regions, including catalytic sites, binding surfaces, and allosteric regions, are strongly constrained during natural evolution. Inverse-folding models have been found to memorize evolutionary constraints [22], thus we ask if they are capturing these non-folding functional constraints to some extent, leading to biased estimates of ΔΔ*G*.

To identify putative functional sites in the natural proteins from the Megascale dataset, we considered sites at which variation caused the largest changes in the protein dark energy [14]. The dark energy *E*^*dark*^ is defined as the difference between the protein sequence free evolutionary energy *E*^*evo*^ and the corresponding physical folding free energy *E*^*fold*^. For a single-site variant it can be computed as

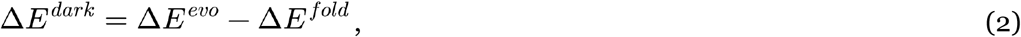

where Δ *E*^*fold*^ correspond to the experimental ΔΔ*G*, if available, and *E*^*evo*^ can be approximated by rescaling the masked logits of the protein Language Model ESM-1v [32] with the folding selection temperature of the protein [14,33] (see *Methods*). Both the evolutionary and folding free energies, and therefore the dark energy, have physical units and can be numerically compared across different protein families. On average, it has been seen that amino acid substitutions at and around annotated functional sites show the largest dark energy loss [14]. To isolate the sequence regions highly constrained by biological functions beyond folding, we sorted the sites of the natural proteins in the Megascale dataset according to the weighted average of *E*^*dark*^ over the 20 possible residues per site, with the natural amino acid frequencies in the dataset as weights.

The average root mean square error (RMSE) between each pure model Δ *E*^*fold*^ and the experimental ΔΔ*G* shows IF models performing better than AWSEM for the complete natural dataset (Fig. 2A). However, IF models are less precise for sites with higher dark energy, while the physical force-field follows the opposite trend. Although IF models have the strongest correlation with the complete deep mutational scanning (DMS) data, their errors increase systematically at sites strongly constrained by biological function beyond folding. At the top functional sites, the experimental data shows only small ΔΔ*G* perturbations, whereas IF models systematically predict larger destabilizing effects (Fig. 2A). In contrast, for those top functional sites, AWSEM predictions are closer to the experimental values than those of any IF model. We also included in our comparison ThermoMPNN [34], a supervised model trained on Megascale ΔΔ*G* labels. Restricting the analysis to its test set, ThermoMPNN exhibits a similar trend and comparable performance to the IF models (Fig. S2).

**Figure 2.**
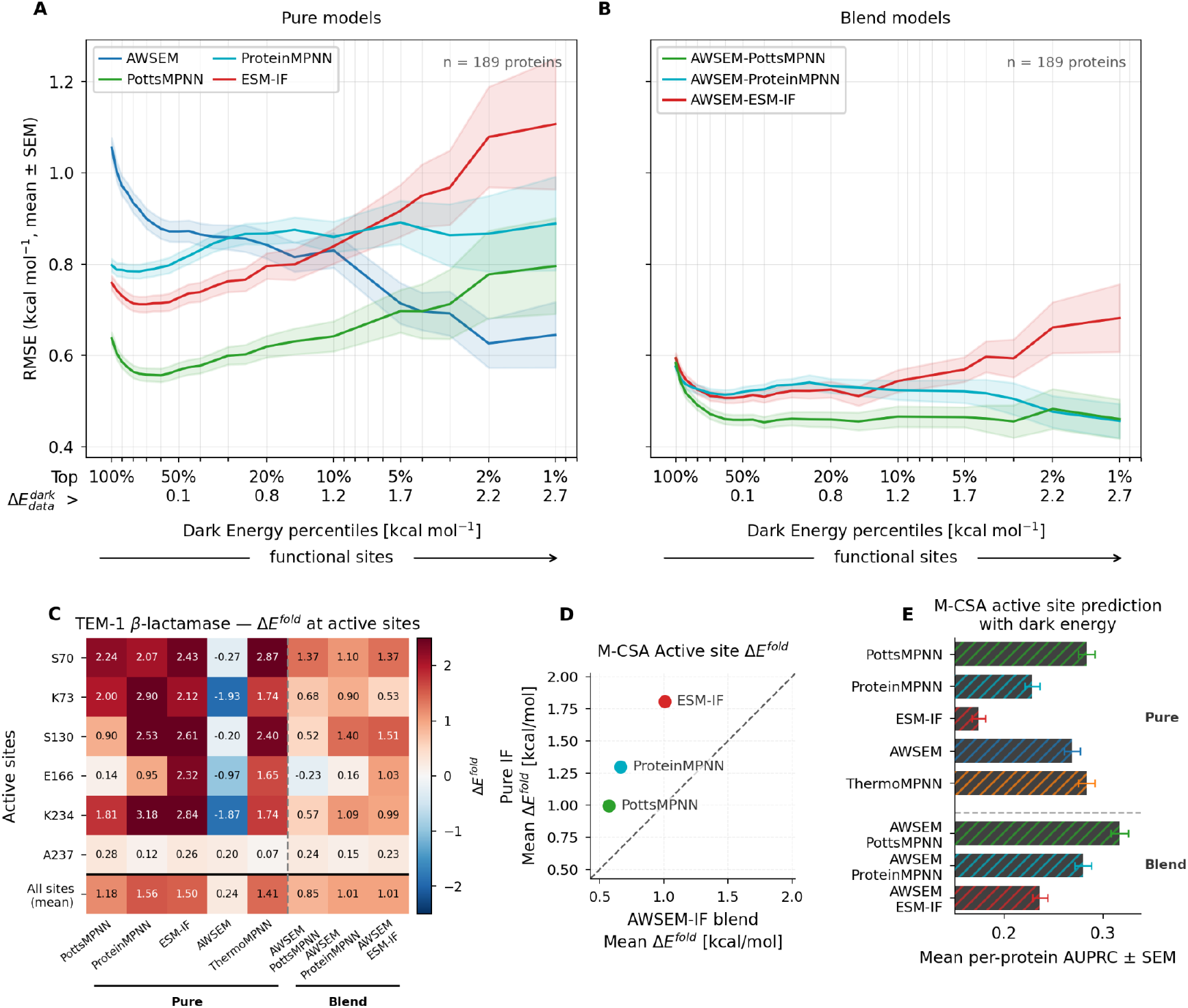
Local discrepancies in folding stability change prediction at functional sites. **A**. Root mean square error (RMSE) between pure model folding energy change predictions and the Megascale experimental values for natural proteins. The RMSE mean and the standard error of the mean (SEM) for each pure model are shown as a function of cumulative percentiles in the site-averaged dark energy, always computed with the experimental data. IF models performed better than AWSEM for all sites, but worse at the top functional sites. **B**. Analogous to panel A, but using the optimal AWSEM-IF blend models. **C**. Weighted-site average folding energy change predicted with each pure and AWSEM-IF model for each annotated active site and on average for all sites for the TEM-1 Beta Lactamase. We include the supervised model ThermoMPNN along with the pure ones. **D**. Folding energy prediction according to pure and blend models for the M-CSA annotated active sites. The average energies are taken over proteins. Error bars (± SEM) are smaller than the symbols and are not visible. **E**. Performance, measured with the AUPRC (Area Under the Precision-Recall Curve), for separating the annotated catalytic sites from the rest for all the enzymes in the Catalytic Site Atlas (CSA) using dark energy computed from different pure and blend folding models. We include the dark energy computed with the supervised model ThermoMPNN.

We repeated the analysis with the AWSEM-IF blend models. We used the optimal blend weights obtained for natural proteins in the correlation analysis (Table S1). The AWSEM-IF blends models show a nearly constant RMSE across the dark-energy percentiles (Fig. 2B). While the pure IF models reach on average RMSE values > 1 kcal/mol at the top functional sites, the AWSEM-ProteinMPNN and AWSEM-PottsMPNN blends maintain RMSE values of approximately 0.4-0.6 kcal/mol across the full range. The blending strategy therefore yields a folding energy predictor that not only improves global correlation with experimental data but also substantially reduces the functional-site bias observed in IF models. An analogous analysis with the Domainome dataset [35] yields equivalent results (Fig. S3A,B).

Instead of using dark energy to identify putative functional sites, for the proteins where both ΔΔ*G* _*bind*_ and ΔΔ*G* _*fold*_ are indirectly measured, we can use ΔΔ*G* _*bind*_ as a proxy for functional constraint. Analysis of KRAS [8], PDZ3, GRB2-SH3 [9] and Src Kinase [10] again show that the more functional a site is, the higher the RMSE of all pure IF models (Fig. S4A), while the blending strategy mitigates the poor performance at the most functional sites (Fig. S4B).

We see a similar behaviour on a dataset where the functional sites are experimentally determined and annotated, the Mechanism and Catalytic Site Atlas (M-CSA) [36]. For the relevant case study of TEM-1 Beta Lactamase, the physics-based force field AWSEM predicts mutations at the active site to stabilize the protein. This is consistent with experimental evidence from beta-lactamases, where removal of the side chain at the catalytic serine increases thermostability while abolishing activity [37], and more broadly with the idea that active-site residues are optimized for catalysis at a cost to protein stability [4,38]. Typically enzyme active sites contain polar, charged, strained, or preorganized groups optimized for substrate recognition, which comes at a cost to the stability of the folded form of the protein. Remarkably, IF models and the supervised model ThermoMPNN show an effect in the opposite direction of the physical force-field, predicting that mutations on the catalytic residues largely destabilize the enzyme (Fig. 2C). In several cases, predictions are more than 1 kcal/mol over the model average for the protein. These results suggest that IF models are biased against the experimentally observed stability-function trade-off, leading them to overestimate Δ *E*^*fold*^ at catalytic residues. Consistently, blending with AWSEM reduces the bias for the case study and across the M-CSA dataset (Fig. 2C,D). Considering all the enzymes studied, on average, IF models predict destabilizations between 1 and 2 kcal/mol, while the blending strategy reduces these values below 1 kcal/mol.

These model discrepancies are also relevant for distinguishing annotated catalytic sites from other sites in the M-CSA dataset. We computed dark energies following Equation 2, using ESM-1v to estimate the free evolutionary energy. We find that blending with AWSEM helps all the IF models to distinguish the catalytic sites from others (Fig. 2E). For the annotated functional sites (Conserved Domain Database (CDD) [39]) at the Domainome dataset [35], IF models predict a much higher folding than dark energy component, whereas AWSEM-IF blends and ThermoMPNN present a more balanced picture (Fig. S3C). In this dataset, the blending strategy only marginally improves the performance of the IF models for separating functional from non functional sites (Fig. S3D).

### Disentangling variant effects characterizes molecular mechanisms of cancer driving genes

We next asked whether AWSEM-IF models can be leveraged to decompose human disease variants into folding-stability and beyond-folding functional components through dark-energy analysis [14]. As a first application, we focused on somatic single-amino-acid substitutions observed in cancer-related genes obtained from the IntOGen platform [40] (see *Methods*). Cancer arises through genetic and epigenetic alterations affecting oncogenes, tumor-suppressor genes and other regulatory programs. Alterations in proto-oncogenes typically drive tumorigenesis by deregulating or hyperactivating their protein products, whereas alterations in tumor-suppressor genes contribute through inactivation mechanisms, often involving reduced activity, abundance, or effective dosage [41–43].

We measured the folding energy change *E*^*fold*^ of the variants with the AWSEM-IF blend model with the best performance on natural proteins, AWSEM-PottsMPNN (λ = 0.6674; Table S1) rescaled to kcal/mol (see *Methods*). For disentangling folding and functional effects, we computed also the dark energy change *E*^*dark*^ produced by the variants using *E*^*fold*^ and ESM-1v to estimate *E*^*evo*^ (equation 2). This gave us a quantitative measure of the effect of each variant on protein folding stability and on other evolutionary constraints, that we interpreted as biological functions beyond folding *E*^*dark*^. The sum of both terms is the free evolutionary energy and measures the total loss of fitness produced by a variant. It corresponds to the ESM-1v scores rescaled to an energetic scale, where the conversion factor is set by the strength of the folding constraints inferred from the evolutionary history of the protein family [14,33]. In the dataset we are using, mutations are enriched for drivers across gene-tumor pairs (see *Methods*), though a residual fraction of passenger mutations is unavoidable. To distinguish likely driver mutations from passengers, we used free evolutionary energy as a proxy for the functional impact of each mutation, reflecting how likely it is to have conferred a selective advantage during tumorigenesis.

For oncogenes and tumor-suppressor genes separately, we compared the folding and dark-energy changes produced by variants at each pathogenicity cumulative percentile (Fig. 3A,B). We found a clear and interpretable dichotomy: in tumor-suppressor contexts, variants are more strongly associated with protein-fold destabilization, whereas in oncogene contexts the deleteriousness signal is mainly carried by the dark-energy component. This separation is more pronounced among the most deleterious variants, for which both the folding-stability loss in tumor suppressors and the dark-energy change in oncogenes reached average values above 2 kcal/mol. This pattern is consistent with the classical distinction between tumor-suppressor inactivation and oncogenic activation introduced above. In this interpretation, the folding component captures loss-of-function effects associated with impaired protein stability or abundance, whereas the dark-energy component captures non-folding effects associated with altered regulatory, catalytic, or interaction mechanisms. Consistently, the average folding and dark energy change per gene generated for the top 10% most pathogenic variants roughly separates oncogenes and tumor suppressors (Fig. 3G). Interestingly, the exception to the rule seems to be IDH1, which we have classified as a tumor suppressor following the TSGene 2.0 database [44], but presents variants with a pure dark energy impact. Indeed, the mutants considered in Fig. 3G on IDH1 are at a single residue, R132, characterized as an gain-of-function hotspot [45]. We report the AWSEM-PottsMPNN folding-stability predictions and the corresponding dark energy changes for all single-site variants in the genes of the studied dataset as Supplementary Data.

**Figure 3.**
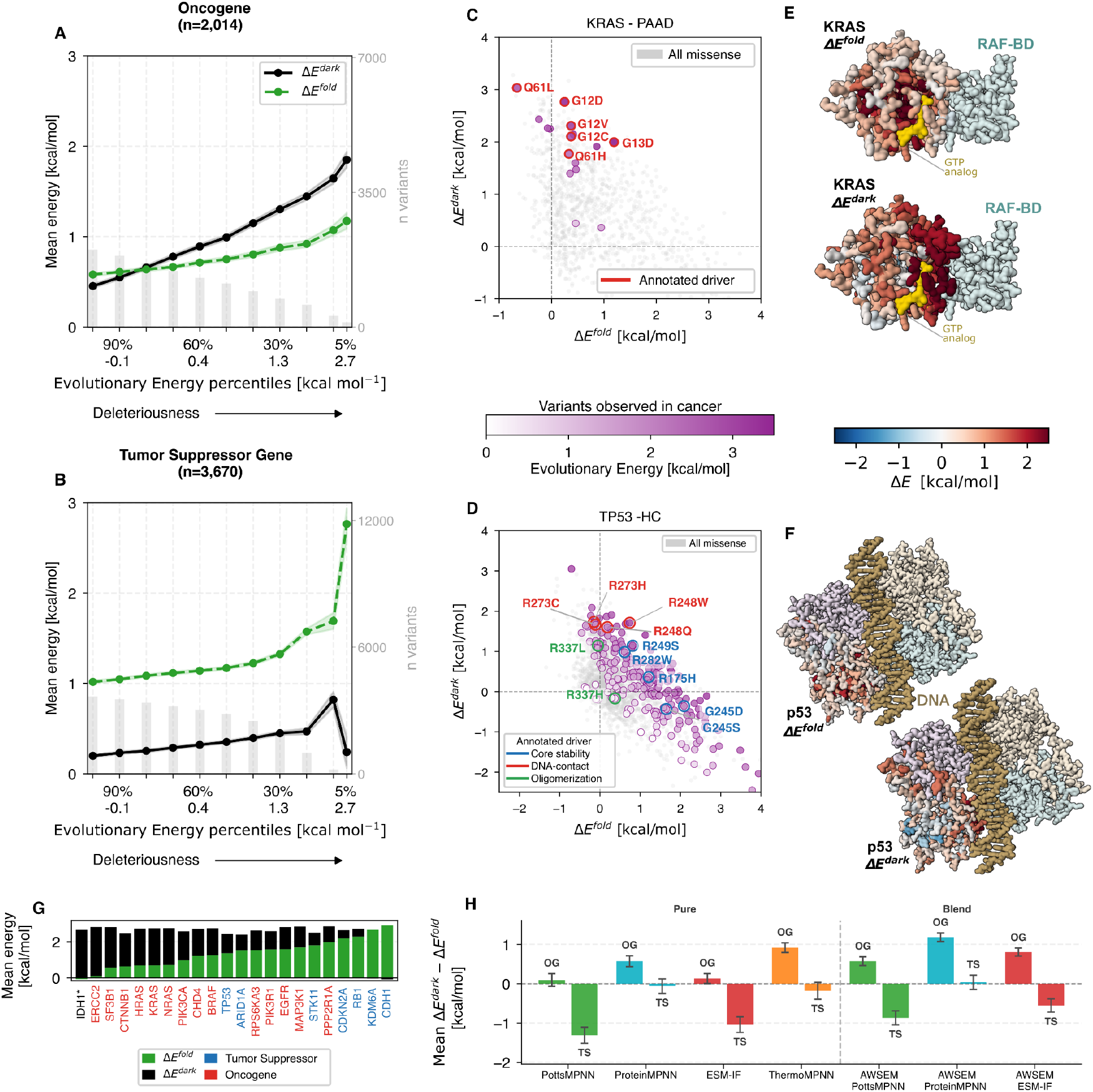
Folding and dark energy of cancer gene variants. **A**. Protein folding-stability (green) and dark (black) energy changes produced by oncogene amino acid variants observed in cancer, as a function of cumulative percentiles of the protein free evolutionary energy, used as a proxy for pathogenicity. The top pathogenic oncogene mutants have a larger impact on biological functions beyond folding (dark energy) than in protein stability. **B**. Same as panel A, for tumor suppressor genes. In this case, variants affect mostly the protein stability. **C**. Dark energy changes as a function of folding energy changes for the oncogene KRAS amino acid variants observed in pancreatic adenocarcinoma. Predicted pathogenicity is shown in a colorscale. Literature annotated drivers are highlighted in red. **D**. Same as panel C, for the tumor suppressor gene TP53 amino acid variants observed in hepatocellular carcinoma. **E**. KRAS structure binding a GTP analog and RAF binding domain, colored by weighted site average folding (top) or dark (bottom) energy changes (PDB: 6vjj). **F**. The structure of p53 tetramer binding to DNA (PDB: 3ts8), where one of the monomers is colored by folding (left) or dark (right) energy changes. A common color scale is provided for all structures in panels E and F. **G**. Folding and dark energy change average per gene for the top 10% most pathogenic variants. Genes are sorted according to the magnitude of the folding energy loss. Oncogenes (red) cluster at the left and tumor suppressor genes (blue) at the right. An exception is IDH1, classified as a tumor suppressor but the studied variants are described as gain-of-function in the literature. **H**. Average difference between the folding and dark energy component for at 10% top pathogenicity for oncogenes (OC) and tumor suppressor genes (TS), according to all studied folding models. This includes the AWSEM-PottsMPNN blend used in all other panels.

We analyzed in detail one case for each class, the oncogene KRAS and the tumor suppressor gene TP53. KRAS encodes a small GTPase that cycles between an inactive GDP-bound and an active GTP-bound state; in the active conformation it recruits effectors including RAF kinases, PI3K and RALGDS to drive cell proliferation. GTP hydrolysis, intrinsically slow in KRAS, is accelerated by GTPase-activating proteins, and oncogenic mutations at positions G12, G13 and Q61 obstruct this hydrolysis, locking KRAS constitutively in its active state. TP53, in contrast, functions as a tetrameric transcription factor that binds specific DNA response elements to activate cell-cycle arrest, senescence and apoptosis, making it the most frequently mutated gene across cancer types [46]. Given this diversity, we focused on one representative cancer context per gene to illustrate the dark-energy decomposition without redundancy: pancreatic adenocarcinoma for KRAS and hepatocellular carcinoma for TP53, two settings where the respective gene is among the most frequently mutated and the driver landscape is well characterized.

We curated literature-based annotations for canonical driver variants, assigning each hotspot to the structural or biochemical mechanism reported in previous experimental and structural studies. For KRAS, driver hotspots were annotated from the Ras mutation survey of Prior, Lewis and Mattos, focusing on the canonical G12, G13 and Q61 variants [47]. For TP53, hotspot variants were annotated using the Joerger–Fersht structure–function classification of common p53 cancer mutants: core-stability/ conformational, DNA-contact and tetramerization-domain mutants [48]. Consistent with these annotations, KRAS driver variants show large dark-energy changes and comparatively small folding-energy changes, clustering in the high-dark-energy/low-folding-energy region of the plot (Fig. 3C), in agreement with their known activating mechanism: they do not primarily unfold KRAS, but instead disrupt the GTPase activity [47]. The KRAS structure colored by dark energy reveals that high dark energy sites concentrate around the nucleotide-binding pocket, switch regions and protein-interaction surfaces (Fig. 3E). TP53 shows a more heterogeneous pattern, consistent with the known diversity of p53 loss-of-function mechanisms: core-stability mutants such as R175H and G245 variants produce stronger protein destabilizing effects, whereas DNA-contact mutants such as R248W and R273H present larger dark-energy changes, and tetramerization-domain variants at R337 form a third class consistent with disruption of p53 oligomerization (Fig. 3D,F).

We compared folding-stability predictions and corresponding dark energy changes to recent experimental deep mutational scanning (DMS) for all single-site variants in KRAS and TP53. For KRAS, Kwon et al. [49] performed gain-of-function and loss-of-function screens in cancer cell lines, and Weng et al. [8] quantified the free energy cost of every KRAS mutation on binding to six interaction partners, including RAF1. Comparison with these assays yields three main messages (Fig. S5). First, the annotated oncogenic driver variants we highlight in Fig. 3C as having high dark energy clearly overlaps with the 86 variants classified as strong gain-of-function by the activation DMS. Second, we find that the dark energy landscape, beside capturing these gain-of-function variants, also captures inactivating variants involved in protein interaction. Through a complementary inactivation screen using a KRAS^G12D^ backbone, Kwon et al. identified two classes of variants decreasing activity: those destabilizing and those disrupting the interaction with other proteins. Weng et al. also identifies those latter variants – concentrated at the S17 in the P-loop, the Switch I triad T35–D38–Y40, and D57 at the Switch II entry point – as disrupting binding to RAF1 without affecting global stability. Indeed, we predict these variants to have large dark energy, highlighting how dark energy aggregates multiple functional signals beyond stability constraints. And finally, third, Kwon et al. showed how variants that mainly affect protein stability map to the secondary-structure elements β4 (residues 77–83), β5/G4 (111–119) and α5 (152–167). In our predictions, variants in these regions carry a large folding energy loss, higher than 3 kcal/mol, while having no dark energy.

For TP53, Funk et al. performed a CRISPR-based saturation genome editing screen [50]. Among the variants identified as loss-of-function by the assay, those in DNA-contact positions show a profile dominated by dark energy. The remaining loss-of-function variants show the opposite pattern, with high folding energy and near-zero dark energy. The dark/folding energy decomposition therefore resolves the two principal TP53 loss-of-function mechanisms: interaction disruption and structural collapse (Fig. S6).

Finally, we compared the average gap between the folding and dark energy component characterizing the top 10% most deleterious variants separated by gene class for the pure and blend models that we studied (Fig. 3H). The blending approach improves the simultaneous prevalence of the folding-stability component for tumor suppressor drivers and the functional dark energy component for oncogene drivers. This highlights the importance of accurately characterising functional sites to reveal the mechanism behind cancer drivers, and shows how combining both IF and physics-based models helps address this challenge.

### Folding and dark energies characterize the molecular mechanism of human disease variants

To evaluate the value of this approach for identifying molecular mechanisms of human disease variants in general, we used a dataset of 714 genes derived from the Human Gene Mutation Database [51], in which single-amino-acid substitutions were classified as gain-of-function (GOF) or loss-of-function (LOF). The LOF class was further subdivided into autosomal recessive (AR), haploinsufficient (HI), and “other LOF” variants. In addition, putatively benign variants from the same genes were obtained from gnomAD and used as neutral controls [7]. For each variant, we computed the folding energy change Δ *E*^*fold*^ using the AWSEM-PottsMPNN blend (Table S1) rescaled to kcal/mol (see *Methods*) and computed the dark-energy change Δ *E*^*dark*^ using the free evolutionary energy from ESM-1v. Predictions for all single-site variants are provided in the Supplementary Data.

We compared folding and dark-energy changes distribution for each mechanism class (Fig. 4A). LOF variants show higher folding-energy changes than dark-energy changes. This suggests that, for LOF variants, pathogenicity is mainly associated with protein destabilization, whereas the purely functional component contributes less. The effect is strongest for HI and AR variants. By contrast, variants labeled as “other LOF,” a class that includes dominant-negative variants, show more balanced folding and dark-energy contributions. We highlight that oligomerization is, in our approach, just another function beyond (monomer) folding. GOF variants are the single class presenting an inverted pattern, with dark-energy changes exceeding folding-energy changes. The control putative benign variants present a dark energy distribution centered near zero, while folding energy distribution is shifted towards more positive values.

**Figure 4.**
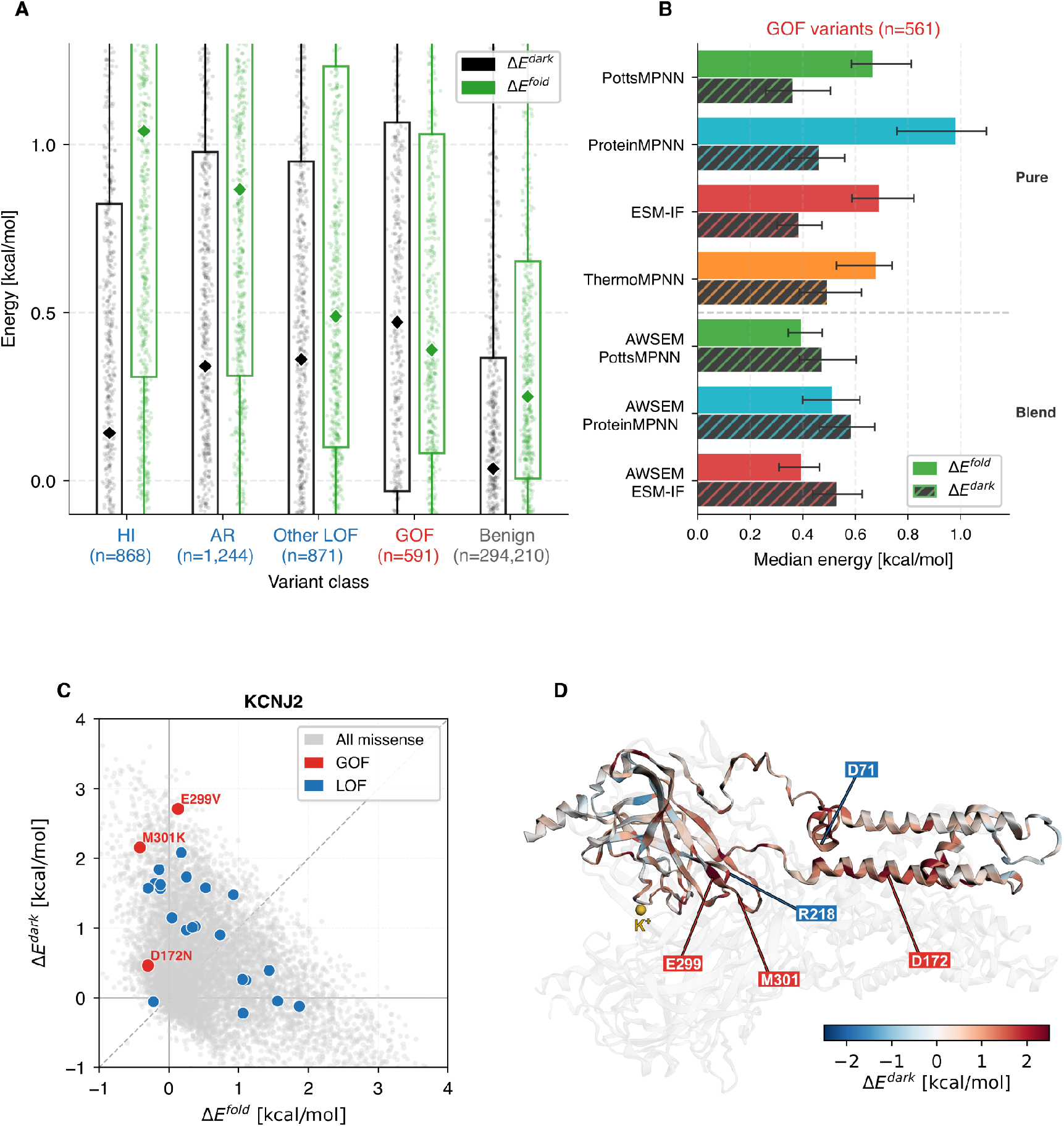
Stability and dark energy of loss-of-function and gain-of-function variants. **A**. Violin plot for the dark and folding energy changes of pathogenic variants annotated as loss-of-function (LOF), including haploinsufficient (HI), autosomal recessive (AR) and other LOF variants and gain-of-function (GOF). Putative benign variants from gnomAD are included as controls. Within the violins, the dots represent the median of the distributions and the lines the interquartile ranges. **B**. Folding and dark energy changes (median) for the gain-of-function variants, according to each pure and blend studied model. Values for the AWSEM-PottsMPNN match panel A. Error bars show 95% bootstrap confidence intervals of the median (2,000 resamples). **C**. Dark energy changes as a function of the folding energy changes for the variants of a case study, KCNJ2, that codifies for the inward rectifier potassium channel Kir2.1. Annotated gain-of-function variants are shown in red and loss-of-function variants in blue. **D**. The tetrameric potassium channel Kir2.1 (PDB: 7zdz). One of the monomers is colored by the weighted site average dark energy changes. Some of the sites with annotated pathogenic variants are labeled, following the color code of panel C.

We analyzed the dark energy and folding-stability loss for gain-of-function variants according to the pure and blend models that we studied (Fig. 4B). Interestingly, all pure models but AWSEM show a larger folding than functional component for GOF variants, including ThermoMPNN. In contrast, AWSEM-IF blends predict a lower destabilization, and therefore a prevalence of the dark energy component.

Among the 60 proteins carrying both GOF and LOF variants in the studied dataset, we analyzed in detail the Kir2.1 inward-rectifier potassium channel, encoded by KCNJ2. Kir2.1 inward rectification is mainly due to voltage-dependent pore block by intracellular polyamines and Mg^2+^, which suppresses outward K^+^ current at depolarized voltages [52]. Fig. 4C shows the folding-energy change predicted with the AWSEM-PottsMPNN blend and the corresponding dark-energy change of Kir2.1 variants. Most annotated variants in our dataset are labeled as loss-of-function, within the “other LOF” subclass. Loss-of-function KCNJ2 variants cause Andersen–Tawil syndrome by reducing Kir2.1-mediated IK1 current; because Kir2.1 channels are tetramers, several experimentally characterized variants act dominantly by suppressing current when co-expressed with wild-type channels [53–55]. Consistent with the mechanistic diversity of this “other LOF” class, these variants affect both folding and dark-energy components. Consistently, several subunit-interface regions show high dark energy (Fig. 4C). In contrast, gain-of-function variants show little predicted folding destabilization, with their pathogenic signal mainly carried by the dark-energy term (Fig. 4B). Consistently, the GOF variants E299V and M301K map to the cytoplasmic pore region controlling inward rectification, which is enriched in dark energy (Fig. 4C). These Kir2.1 GOF variants have been shown experimentally to increase outward IK1 through reduced inward rectification and are associated with short QT syndrome phenotypes [56,57].

## Discussion

Natural selection shapes protein sequences under overlapping and sometimes contradictory folding-stability and functional constraints [3]. Variant effect predictors trained on evolutionary data, as single readout functional assays, are blind to this distinction [58–60]. For human missense variants, drawing this distinction could improve diagnosis and enable new therapies, for instance by pharmacological rescue [61–63]. Going beyond one-dimensional deleteriousness or pathogenicity prediction, therefore requires predicting folding free energy changes (ΔΔ*G*) at high throughput and disentangle them from other functional components. A fast, scalable, and unsupervised alternative to experimentally estimate ΔΔ*G* is to use Inverse Folding (IF) models, trained on structure-to-sequence correspondence. Our results show that IF models, such as ESM-IF1, ProteinMPNN and PottsMPNN, sometimes misread a purely functional alteration as a loss of stability, highlighting an issue for cleanly separating folding-stability contributions, especially at sites that are functionally relevant. However, blending them with the physics-based model AWSEM, both improves the global correlation with experimental data (Fig. 1) and reduces IF bias at functional sites (Fig. 2). The improvement goes beyond simple noise reduction from averaging; it reflects the combination of two complementary approaches. AWSEM is a transferable coarse-grained potential grounded in the energy landscape theory of protein folding, and unlike IF models, its burial and contact parameters are optimized to maximize the stability gap between the native state and an ensemble of misfolded decoys, without relying on evolutionary sequence signals [26,64].

Interestingly, AWSEM performed better than any IF model for the Megascale de novo proteins (Fig. 1D). This drop in IF performance on an out-of-distribution task could reflect overfitting to naturally evolved sequence-to-structure patterns. On natural proteins, AWSEM avoids the systematic bias at functional sites shown by the IF models and by the supervised model ThermoMPNN (Fig. 2, Fig. S2, Fig. S3, Fig. S4). These inverse-folding model distortions are evident, for instance, at the TEM1 β-lactamase catalytic residues, where IF models systematically predict mutations at active sites to be largely destabilizing, at least in part contradicting experimental evidence [37] and physics-based computational analysis [27,65,66]. Consistent with this, some functional residues in the β-lactamase fold are known to resist stability optimization by IF models [67]. By conflating evolutionary conservation with folding-stability requirements, IF models misinterpret some functional constraints as structural ones. The AWSEM-IF blends combine the strengths of both approaches, yielding a family of unsupervised models suited for probing folding-stability changes with improved global accuracy and reduced biases at functional sites.

Using the best performing AWSEM-PottsMPNN blend on natural proteins and ESM-1v [32] to estimate the evolutionary energy, we disentangled the effect of human missense variants into folding-stability and dark energy components. Although log-likelihoods from protein language models have known limitations as fitness estimates [68–70], we take ESM-1v as an approximate evolutionary energy leaving improvements for future work.

Somatic variants in oncogenes and tumor suppressor genes follow quantitatively distinct biophysical trajectories (Fig. 3A,B). Oncogenic drivers mostly preserve stability, but perturb functions beyond folding, reaching almost 2 kcal/mol of dark-energy loss on average, twice the threshold associated with significant functional perturbation in protein domains [14] and pointing to strong disruption of catalytic or regulatory function. For instance, gain-of-function KRAS mutants that lock the enzyme in its active GTP-bound state are high dark-energy variants that leave the folding-stability largely unchanged (Fig. 3C,E, Fig. S5). This is consistent with previous structural characterizations of oncogenes as difficult to distinguish from non-cancer genes except for the clustering of drivers in active sites [71]. In contrast, strongly pathogenic somatic variants in tumor suppressor genes produce an average folding-energy loss over 2-3 kcal/mol (Fig. 3B), matching the well-described inactivation of tumor suppressors through structural destabilization [72]. We further disentangle the functional diversity of TP53 mutational landscape, recovering the annotated core-stability, oligomerization related and DNA-contact mutants [48], without any biochemical input (Fig. 3D), yielding a mechanistic characterization of every variant in silico and without supervision (Fig. S6). This is directly relevant therapeutically: pharmacological rescue approaches targeting p53 have been guided precisely by the distinction between stability-impaired and DNA-contact mutants [48,61,73]. More broadly, this approach opens a promising avenue for the characterization of unknown molecular mechanisms underlying cancer and driver gene discovery.

The folding/dark energy spectrum extends beyond cancer to germline pathogenic variants with annotated gain- and loss-of-function effects. For the inward-rectifying potassium channel Kir2.1, encoded by KCNJ2, predictions of folding and dark energy changes (Fig. 4) separate gain-of-function mutants associated with short QT syndrome phenotypes [56,57] from the channel inactivating mutants causing Andersen–Tawil syndrome [53–55]. Beyond this case study, gain-of-function variants show, on average, a higher dark than folding-energy loss (Fig. 4A). In contrast, haploinsufficient and autosomal recessive variants are predominantly destabilizing, compatible with structure-based studies [7], while dominant-negative variants show more balanced contributions from both components, consistent with their oligomeric mechanism of action.

We have presented a blending strategy that exploits the complementarity between physics-based and deep-learning approaches to improve folding-stability change prediction. AWSEM-IF blends improve the already strong performance of inverse-folding models on site-saturation data, gaining sensitivity to the stability-function trade-off in protein molecules captured by AWSEM. Although we tested a limited number of models, the theoretical basis of the blending means it generalises naturally to other IF models and physics-based potentials at different resolutions, already successful in other hybrid approaches [74]. Direct applications of the AWSEM-IF blend include protein design through stability enhancement. Recent observations that functional sites resist stability optimization in backbone-conditioned sequence generation [67] suggest that de novo design may also benefit from AWSEM-IF blends, in which the physical force-field corrects for the stability-function conflation inherent to IF models.

The framework presented here also has important limitations. The current implementation of AWSEM-IF blends has only been applied to single-site variants, leaving epistatic and multi-site effects out of scope. And while dark energy is the best available estimator of the energetic cost of protein function beyond folding, is not necessarily a physicochemical functional energy in a strict sense: its thermodynamic interpretation is clearest when the selection temperature for function equals that for folding, as in binding, but the scale may differ for other chemical activities such as catalysis. These limitations notwithstanding, the results presented here suggest a broader principle: the divergence between the evolutionary and the folding energy landscapes, quantified here as dark energy, is not noise to be minimized but information to be interpreted. As new folding-stability models trained on sequence and structure evolutionary information become available, blending strategies of the kind introduced here offer a path toward variant effect predictors that are not only accurate but mechanistically transparent.

## Methods

### Folding Stability Experimental Data

We used 335 deep mutational scanning assays of the “Mega-scale experimental analysis of protein folding stability” [31]. These included the assays for 189 natural protein domains and the 146 de novo designs of the database. From the complete dataset, we only excluded the ones where the assayed background already carries one or more mutations. We used the thermodynamic folding stability measurements (ΔΔG) for single-site amino acid substitutions, imposing the sign convention where ΔΔG < 0 for stabilizing variants, and the corresponding AlphaFold2 model structures.

### Reference Protein Dataset

To compute the folding energy change standard deviation for each model, we used as a reference dataset a set of 200 human proteins randomly obtained from the AlphaFold Protein Structure Database (https://alphafold.ebi.ac.uk/) [75]. We used the provided sequences and the structure predictions generated using AlphaFold2 [76].

### Enzyme Active Site Data

We studied the PDB for all the entries of the Mechanism and Catalytic Site Atlas (M-CSA) [36]. After PDB cleaning and sequence matching with the annotations, we considered 785 enzymes, including 263 monomers and 522 multimers.

### Somatic Mutation Data

Cancer somatic mutation data were obtained from the IntOGen platform [40], which integrates whole-exome and whole-genome sequencing cohorts from the Cancer Genome Atlas (TCGA), the Pan-Cancer Analysis of Whole Genomes (PCAWG) project, the International Cancer Genome Consortium (ICGC), and cBioPortal. Following the filtering procedure of Muiños et al. [77], we restricted the analysis to somatic mutations, defined as those present in the tumor but absent in the germline, and retained only gene-tumor pairs with an estimated driver mutation fraction of 0.85 or higher, as determined by dNdScv [78], which estimates gene-specific non-synonymous to synonymous substitution ratios (dN/dS) corrected for regional variability in neutral mutation rate, consequence type, and tumor-specific mutational processes. Gene-tumor pairs with fewer than 30 observed mutations were additionally excluded. This yielded 249 gene-tissue combinations for downstream analysis. Genes present in the TSGene 2.0 database [44] were classified as tumor suppressors; all remaining driver genes were classified as oncogenes. Although these filtering steps ensure that the vast majority of retained mutations are likely to be drivers, a residual fraction of passenger mutations cannot be entirely excluded.

### Loss-of-function and gain-of-function variants dataset

We used a dataset of 714 genes with variant-level annotations derived from the Human Gene Mutation Database using natural language processing (NLP) by Bayrak et al. [51] and curated afterwards by Gerasimavicius et al. [7]. The correspondent predicted structures were obtained from the AlphaFold Protein Structure Database (https://alphafold.ebi.ac.uk/) [75].

### AWSEM

We used the transferable Associative Memory Water Mediated, Structure and Energy Model (AWSEM) coarse-grained force field [26]. The folding energy change for a single-site variant 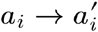 is computed as 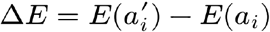, evaluating only the terms that depend on residue *i* and its contacts, with the backbone coordinates held fixed at the native structure. Two terms of the potential were considered, the burial and the contact terms, following the force-field usage for dark energy [14] and frustration analysis [24,27]. The burial term is a many body interaction which is based on a particular residue type’s propensity to be in a low density environment. The contact term distinguishes three contact types: direct, and water- or protein-mediated.

Calculations were performed with the AWSEM implementation in the Frustratometer Python package available at https://github.com/HanaJaafari/Frustratometer/tree/galpern2025 [79]. In this implementation, the AWSEM energy function can be written in a Potts model form, with fields *h*_*i*_ (*a*_*i*_) corresponding to the burial term and couplings *J*_*ij*_ (*a*_*i*_, *a*_*j*_) to the contact term, allowing efficient computation of single-site energy differences. In this work, sequence pairs within | *i* − *j* | < 3 were excluded from the density calculation and pairs within | *i* − *j* | < 2 from the contact energy.

### Inverse Folding Models

We used the unsupervised models ESM-IF1 [17], ProteinMPNN [18] and PottsMPNN [21]. These models assign a score to an amino-acid sequence given a fixed protein backbone. For each structure *S* and single-site mutation *s*_*i*_ = *a* → *b*, we scored the change in model preference between the mutant and wild-type residue at the same position. We used the sign convention that larger scores correspond to mutations that are less compatible with the native structure. For ESM-IF1, to compute the log-probability difference between the mutant and wild-type amino acid at position *i* from the structure-conditioned residue probabilities,

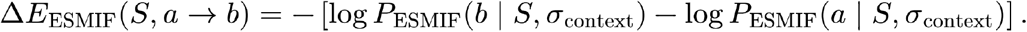

Here, σ_context_ denotes the sequence context used by the autoregressive inverse folding model. Thus, substitutions assigned lower probability than the wild-type residue receive positive scores. For long proteins, ESM-IF1 was evaluated using overlapping structural chunks. Proteins longer than 500 residues were split into windows of at most 500 residues with 64-residue overlaps. Each target window was scored as chain A, while additional chains provided structural context from the remaining parts of the protein. Scores in overlapping regions were merged across windows. This allowed us to score proteins longer than the ESM-IF1 practical single-chain limit.

For ProteinMPNN, we similarly computed the change in model preference for the mutant residue relative to the wild-type residue at each position,

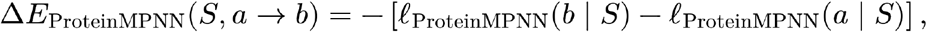

where *ℓ*_ProteinMPNN_(*x* | *S*) is the ProteinMPNN logit for amino acid *x* at the mutated position. We scored all possible single amino-acid substitutions for the selected chain of each structure. ProteinMPNN uses local structural neighborhoods and was run directly on the input structures without using a chunking procedure.

For PottsMPNN, we used the model in single-mutant scanning mode to compute the predicted mutational folding score for each substitution,

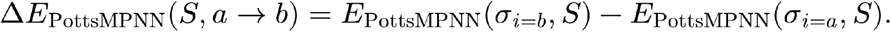

The PottsMPNN output was used in the mutant-minus-wild-type direction, so larger values indicate mutations predicted to be more destabilizing or less compatible with the native backbone.

### ThermoMPNN

We also evaluated ThermoMPNN [34], a supervised model trained on Megascale ΔΔG labels to predict mutational stability changes. For each input structure, ThermoMPNN was run in single-site saturation mutagenesis mode. For every residue position and every amino-acid substitution, the model predicts a mutational stability change. The ThermoMPNN score was used directly in the mutant-minus-wild-type direction, where larger positive values indicate mutations predicted to be more destabilizing.

### Folding Energy Units

To compare the folding energy Δ *E*^*fold*^ values obtained with models and the experimental ΔΔ*G*, we rescaled model results to kcal/mol following Δ *E*^*fold*^ [kcal / mol] = Δ *E*^*fold*^ *σ ΔΔG / σ Δ E*^*fold*^, as has been previously done [14]. Here σ Δ *E*^*fold*^ is the representative standard deviation of a model, computed as the average over the protein DMS of a representative dataset, in this case a reference dataset of 200 human proteins (see above). Results are shown at Table S2. The representative standard deviation for the experimental ΔΔ*G* was computed as the average over the Megascale dataset natural proteins, finding a value σ ΔΔ*G* = 0.9777 of kcal/mol.

### Large Protein Handling

The AlphaFold Protein Structure Database provides in the case of proteins longer than 2700 amino acids (aa) shorter overlapping fragments of 1400aa [75]. We leveraged this separation of large structures to run each folding model over each fragment separately. We afterwards combined the scores, taking the average for overlapping regions.

### Free Evolutionary Energy

We estimate the changes in the free evolutionary energy rescaling for each protein the dimensionless Massieu potential or evolutionary score Ψ^*evo*^ with the correspondent folding selection temperature 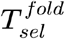 [14],

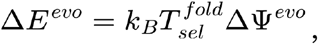

where *k*_*B*_ is the Boltzmann constant.

### Evolutionary Score

To compute the changes in the evolutionary score upon single-site variants, ΔΨ^*evo*^, we computed the masked marginal difference with the ESM1v model (650M parameters, 33 layers) as in [32]. For a given protein sequence *σ* = (*σ*_1_, … *σ*_*L*_), we take the evolutionary score for each possible single-residue substitution. This is computed with a forward pass for each position by masking that position then taking the log softmax function of the output logits in that position, *l*_*i*_ = log *P* (*σ*_*i*_ | *σ*_\*i*_). For a mutation *σ*_*i*_ = *a* → *b*, the evolutionary score is a log-likelihood ratio between the mutant and the wildtype residues,

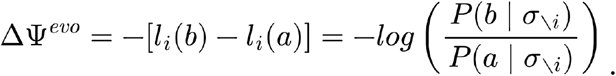

We choose the sign convention such that higher scores indicate mutation is less likely under the model. Because ESM-1v accepts a maximum input length of 1024 tokens, long proteins were scored using local windows containing up to 1022 amino-acid residues, allowing for start and end tokens where relevant. These windows were centred around the amino acid position being masked and scored in each forward pass.

### Folding Selection Temperature

We estimated the apparent temperature at which sequences were selected by evolution for a particular protein family or fold [2,33]. In practice, 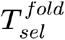 is used in this work as a protein-specific scaling factor to make the evolutionary score Ψ^*evo*^ comparable to the folding free energy changes ΔΔ*G*, or equivalently the corresponding folding model predictions Δ *E*^*fold*^. Following [14,30], such factor is computed as the ratio of the two standard deviations of the folding free energy and the evolutionary score changes over all single-site variants of the protein, 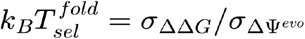, where *k*_*B*_ is the Boltzmann constant.

### Energy weighted site averages

All the site-wise folding and dark energy values in this work were computed by taking a weighted average per site using as weights natural amino acid frequencies. For human protein datasets, we use the frequencies obtained from the human proteome subset. For the Megascale and the enzyme set (M-CSA), we used their own amino acid abundance for the weights.

## Code and data availability

The code is available at https://github.com/DiasFrazerGroup/blend_and_disentangle. Supplementary data, including precomputed scores for all datasets and to reproduce the main figures are available at https://doi.org/10.5281/zenodo.21241135

## Acknowledgments

We thank other members of the Dias and Frazer lab for their thoughtful feedback and many interesting discussions throughout the development of this work. We acknowledge the EuroHPC Joint Undertaking for awarding us access to MareNostrum5 at BSC (project EHPC-DEV-2026D01-121). E.A.G. acknowledges support from the European Union’s Horizon Europe under the grant agreement No 101208464. E.A.G. also acknowledges L.A.M.C. for his recent inspiring performance. X.S.S acknowledges the support of the Spanish Ministry of Science and Innovation (PRE2022-102503 funded by MICIU/AEI /10.13039/501100011033 and ESF+). F.B. acknowledges the support of the European Union’s Horizon 2020 research and innovation programme under the Marie Skłodowska-Curie grant agreement No. 713673 (fellowship code LCF/BQ/DI23/11990061). This work was supported by the Spanish Ministry of Science and Innovation (PID2022-140793NA-I00) funded by MCIN /AEI /10.13039/501100011033 / FEDER, UE). We acknowledge support of the Spanish Ministry of Science and Innovation through the Centro de Excelencia Severo Ochoa (CEX2020-001049-S, MCIN/AEI /10.13039/501100011033), and the Generalitat de Catalunya through the CERCA programme. Research for this publication has been partially carried out in the Barcelona Collaboratorium for Modelling and Predictive Biology.

Funded by the European Union. Views and opinions expressed are however those of the author(s) only and do not necessarily reflect those of the European Union. Neither the European Union nor the granting authority can be held responsible for them.

## Supplementary Information

### Supplementary Methods

#### Inverse-folding background corrections

We tested the single-residue background corrections to ESM-IF1 and ProteinMPNN from Dutton et al. [20]. Following the authors, for ESM-IF1, we also included the Metheonine correction. These corrections were used only in the explicitly labeled corrected inverse-folding analyses, such as Fig. S1.

#### Analysis on the Human Domainome dataset

We used the Domainome dataset [35], a set of human protein domains where functional sites are annotated according to the Conserved Domain Database (CDD) [39]. The full dataset comprises 522 domains spanning 97 PFAM families, covering 30,124 residue positions, of which 2,818 (9.4%) are annotated as functional by CDD. We used the AlphaFold predictions for the protein structures provided by the Authors [35]. For the RMSE analysis shown in Fig. S3, we used the Domainome experimental fitness measurements as a ΔΔ*G*-like reference after sign inversion, such that lower fitness corresponds to larger deleterious effect. To place this reference on the same scale used for the model energies, inverted fitness values were multiplied by the average natural megascale experimental ΔΔ*G* standard deviation and divided by the mean, across Domainome proteins, of the per-protein standard deviation of inverted fitness. Sites were then ranked by the experimental dark-energy computed using the rescaled inverted fitness, and RMSE was evaluated for cumulative top-percentile thresholds of this score. To evaluate the ability of each model to prioritize functional sites, we computed the Area Under the Precision-Recall Curve (AUPRC) per domain, treating CDD-annotated residues as positives and all remaining residues as negatives. Domains for which no residue carried a CDD annotation were excluded from the analysis, as AUPRC is undefined in the absence of positive examples; this criterion removed 276 of 522 domains. The remaining 246 domains, distributed across 44 PFAM families, were retained for evaluation. To avoid inflating performance estimates through families represented by many homologs, we first averaged the per-domain AUPRC within each PFAM family, then computed the mean and standard error of the mean (SEM) of those 44 family-level averages. This two-level aggregation is reported in Fig. S3D.

#### Analysis on Double Deep Mutational Scannings

We considered the dual-readout DMS data from Lehner’s Lab for KRAS [8], PDZ3, GRB2-SH3 [9] and Src Kinase [10]. We used the values for single-site variants of ΔΔ*G*_*bind*_ (or ΔΔ*G* _*activity*_ in the case of Src Kinase) and ΔΔ*G*_*fold*_ obtained by the authors with the MoCHI model. The correspondent predicted structures were obtained from the AlphaFold Protein Structure Database [75]. For the folding models performance at functional site analysis (Fig. S4) we used ΔΔ*G*_*fold*_ as the reference to compute each model RMSE and we calculated the cumulative percentiles for the weighted site average of ΔΔ*G*_*fold*_ (or ΔΔ*G*_*activity*_).

## Supplementary Tables

**Table S1.** Optimal weights for the AWSEM-IF blend models. For each subset of the Megascale dataset and for each AWSEM-IF blend model, the maximum value of Pearson’s r and the corresponding blend weight.

| Data subset | N | Model 1 | Model 2 | Max Pearson's r | $\lambda$ |
| --- | --- | --- | --- | --- | --- |
| natural | 189 | AWSEM | PottsMPNN | 0.7377 | 0.6674 |
| natural | 189 | AWSEM | ProteinMPNN | 0.7194 | 0.5856 |
| natural | 189 | AWSEM | ESM-IF1 | 0.7242 | 0.6078 |
| de novo | 146 | AWSEM | PottsMPNN | 0.6945 | 0.4815 |
| de novo | 146 | AWSEM | ProteinMPNN | 0.6720 | 0.3689 |
| de novo | 146 | AWSEM | ESM-IF1 | 0.6947 | 0.4579 |
| all | 335 | AWSEM | PottsMPNN | 0.7189 | 0.5864 |
| all | 335 | AWSEM | ProteinMPNN | 0.6987 | 0.4912 |
| all | 335 | AWSEM | ESM-IF1 | 0.7114 | 0.5425 |

**Table S2.** Average standard deviations for each model folding energy change over DMS of the Megascale natural proteins.

| **Model** | $\sigma_{\Delta E}$ |
| --- | --- |
| AWSEM | 12,743 |
| PottsMPNN | 3,441 |
| ESM-IF1 | 5,163 |
| ProteinMPNN | 2,006 |
| ThermoMPNN | 0,977 |

## Supplementary Figures

**Figure S1.**
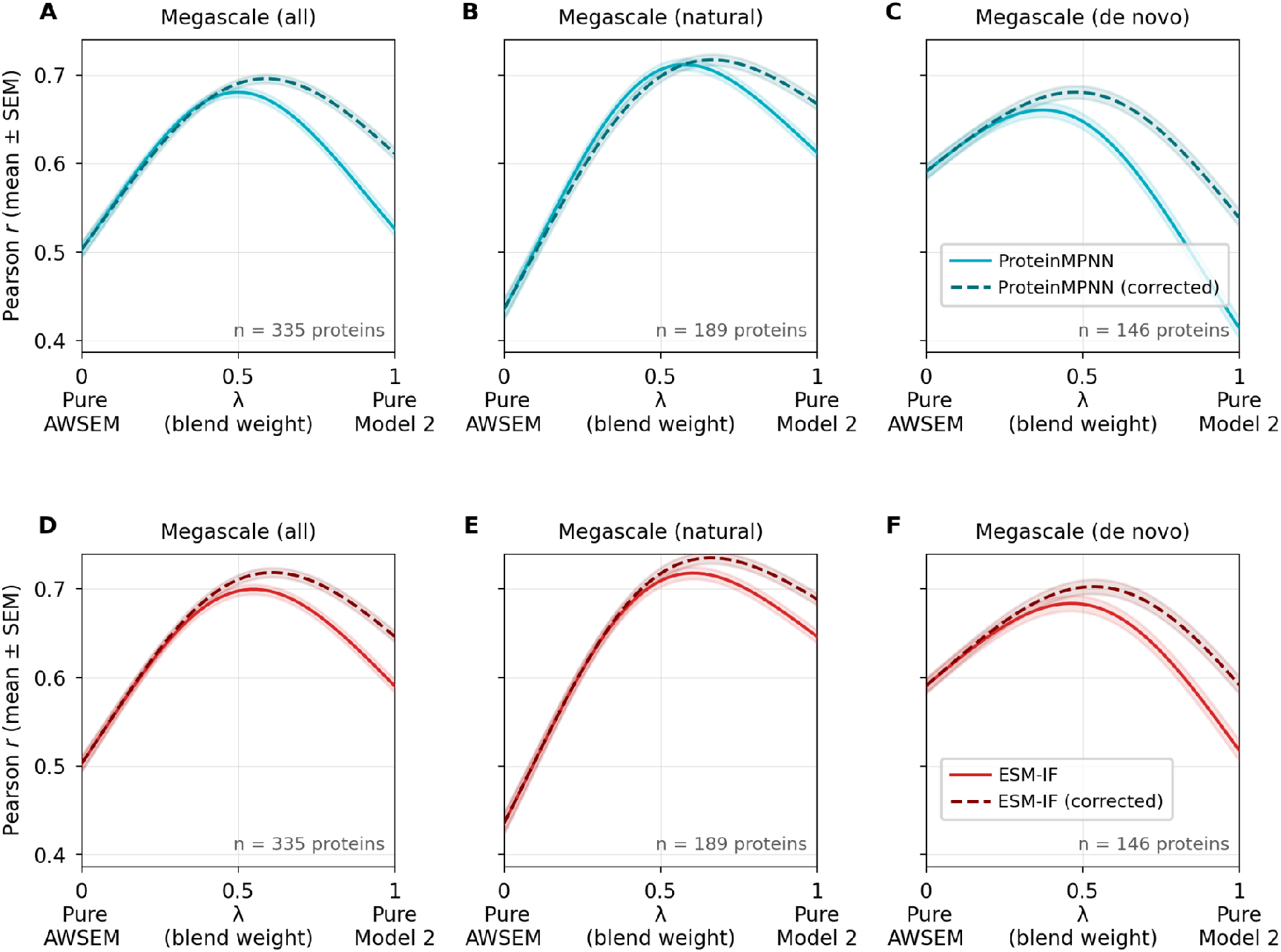
Performance of AWSEM-IF blends for the corrected ProteinMPNN and ESM-IF models. **A**. Correlation between AWSEM-ProteinMPNN blend model predictions with and without single-residue background corrections (Dutton et al. [20]) and the Megascale experimental data measured with the Pearson’s r coefficient mean over all single-site variants of natural and de novo proteins and its standard error (SEM), as a function of the blend weight. **B**. Same as panel A, for only the natural proteins in the dataset. **C**. Same as panel A and B, for only the de novo designs in the dataset. **D-F**. Same as panels A-C for AWSEM-ESM-IF blend model predictions with and without single-residue background corrections.

**Figure S2.**
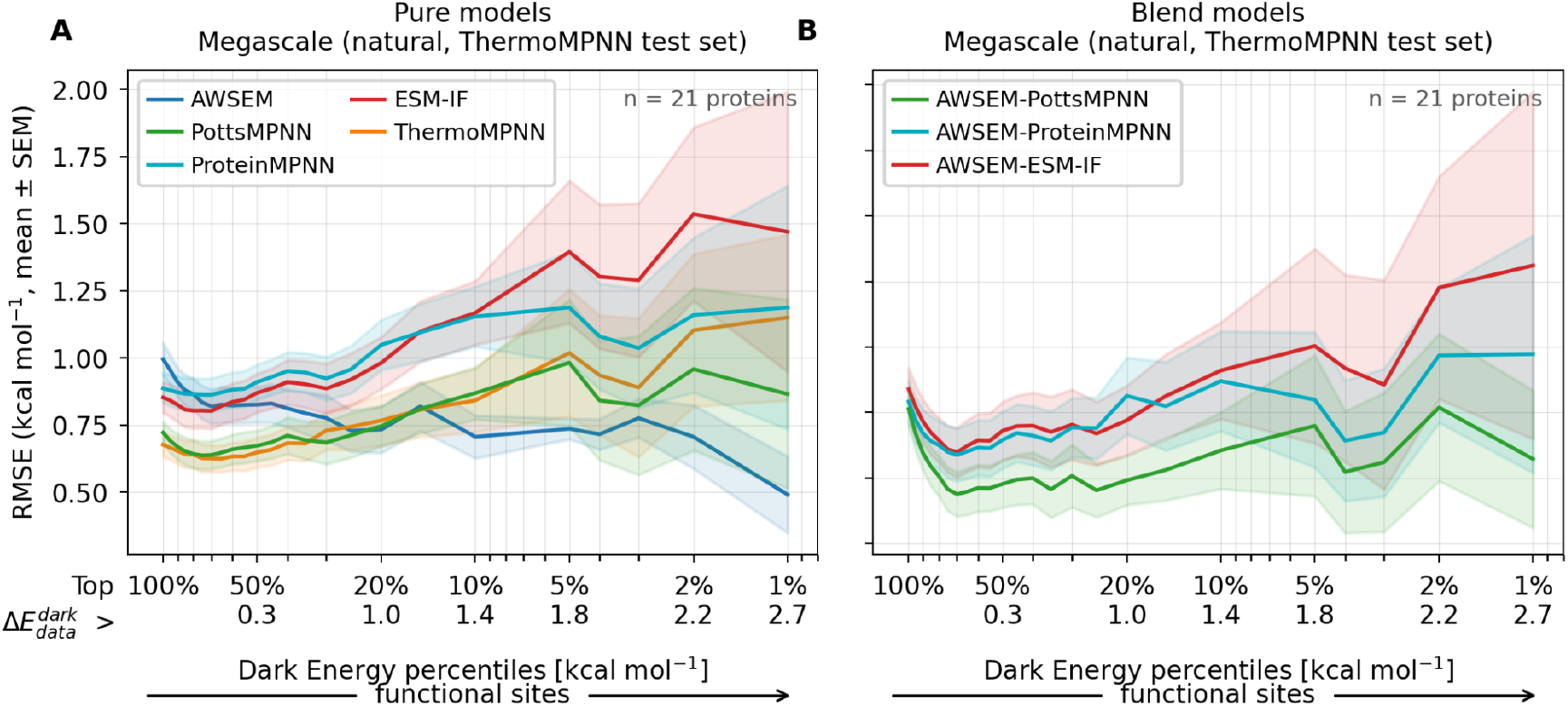
Model performance including ThermoMPNN for Megascale natural proteins. **A**. Root mean square error (RMSE) between pure model folding energy change predictions and the Megascale experimental values for natural proteins. To evaluate the supervised model ThermoMPNN (orange line),the protein set is restricted to the test set of ThermoMPNN. The RMSE mean and the standard error of the mean (SEM) for each pure model are shown as a function of percentiles in the site-averaged dark energy, always computed with the experimental data. **B**. For the same protein set, RMSE of the blend models.

**Figure S3.**
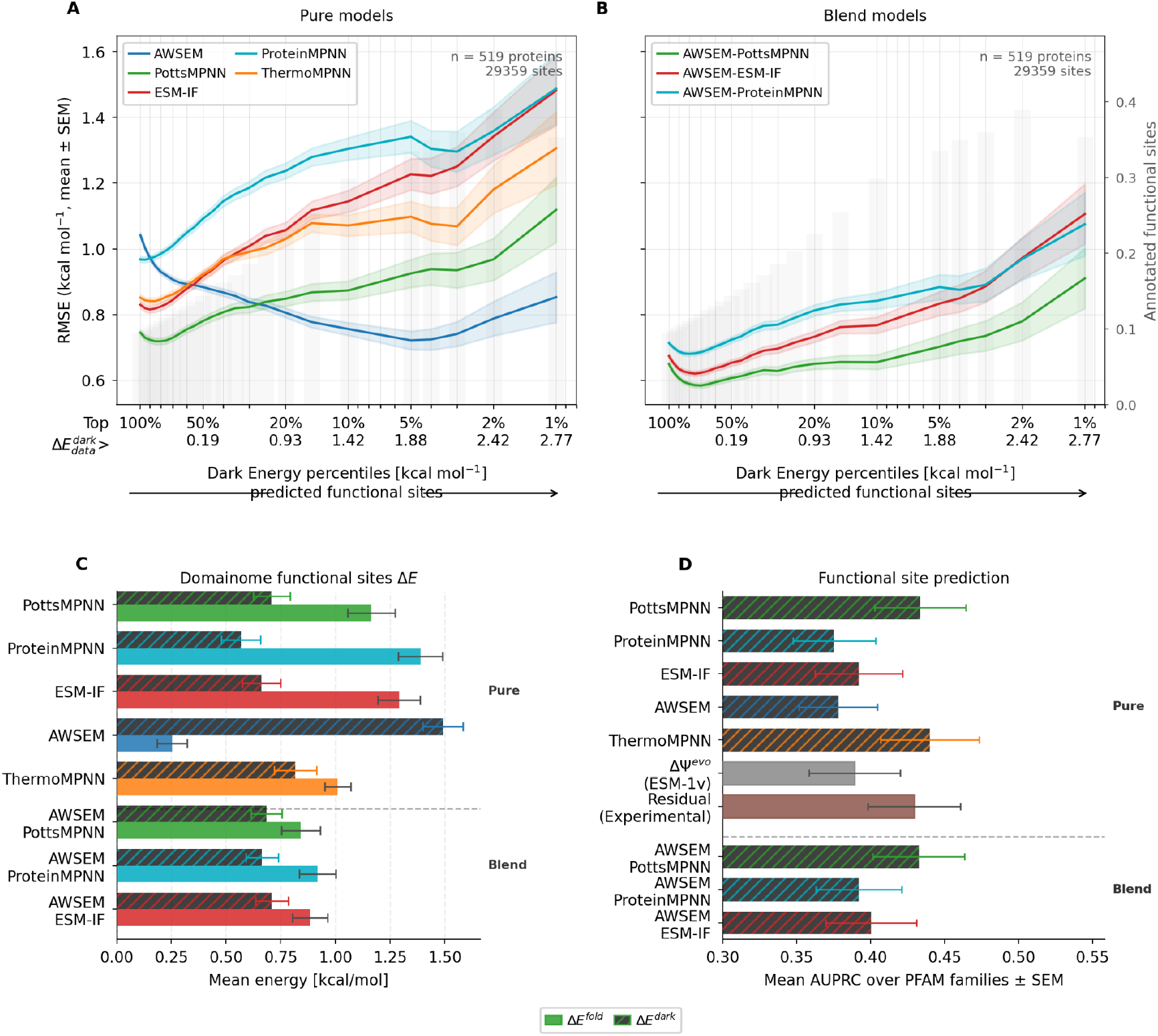
Model performance at functional sites for Domainome data. **A**. Root mean square error (RMSE) between pure model folding energy change predictions and rescaled experimental fitness (experimental proxy for ΔΔG) for the Domainome dataset. The RMSE mean and the standard error of the mean (SEM) for each pure model are shown as a function of percentiles in the site-averaged dark energy, always computed with the experimental data. The fraction of annotated functional sites (CDD) per percentile is shown at the background as a barplot. **B**. Analogous to panel A, but using the optimal AWSEM-IF blend models. **C**. Folding and Dark Energy changes for the CDD annotated functional sites in the Domainome dataset. The average energies are taken over protein and then over PFAM protein families and are shown with the corresponding standard error. **D**. Performance for separating the CDD annotated functional sites from the rest for the human protein domains of the Domainome dataset. The average of the AUPRC over protein and then over PFAM protein families is shown with the corresponding standard error.

**Figure S4.**
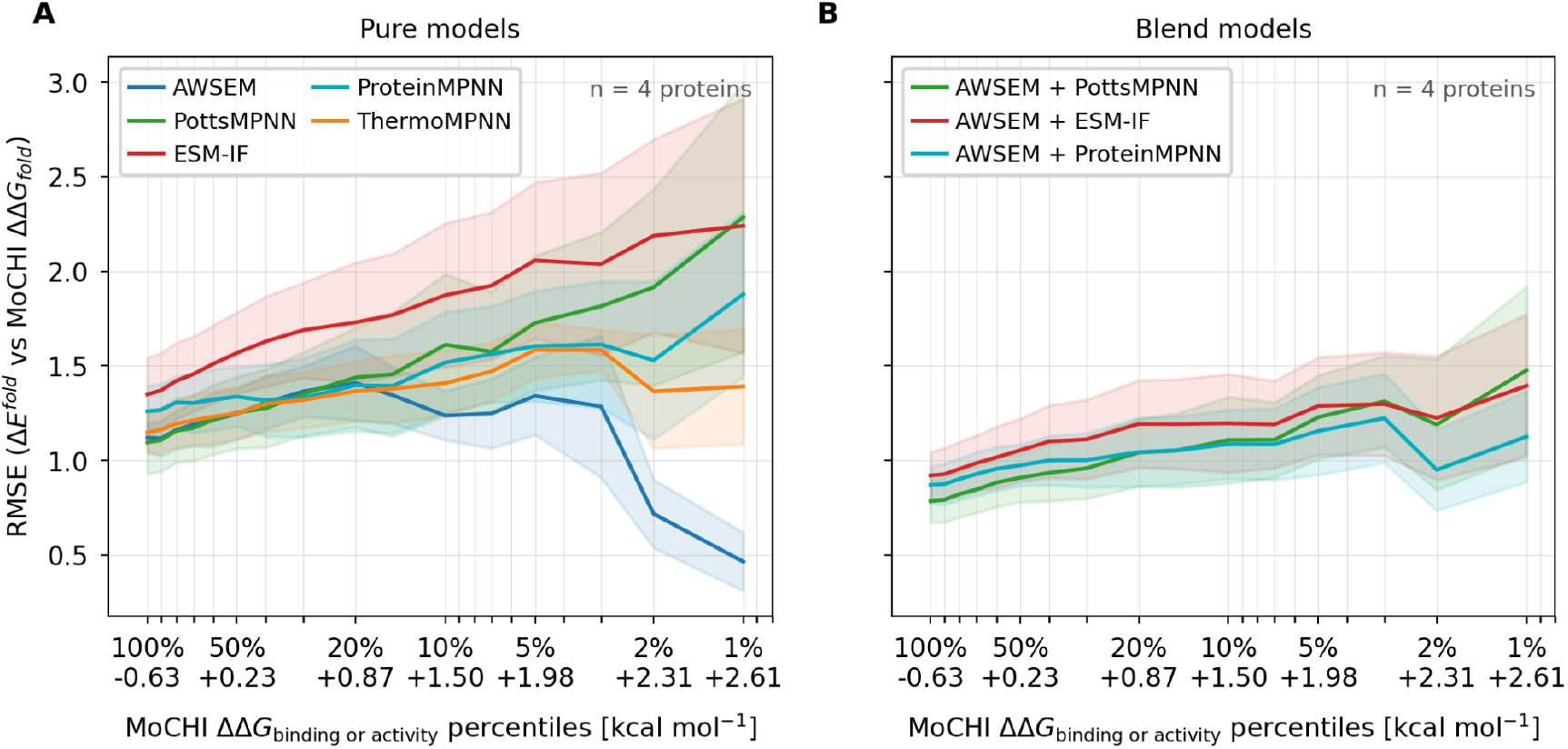
Model performance at functional sites for the double DMS data. **A**. Root mean square error (RMSE) between pure model folding energy change predictions and ΔΔG_folding_ computed at Lehner’s Lab with the MoCHI model from dual-readout DMS data for KRAS, PDZ3, GRB2-SH3 and Src Kinase. The RMSE mean and the standard error of the mean (SEM) for each pure model are shown as a function of percentiles in the site-averaged ΔΔG_binding/activity_, also computed from the dual-readout DMS data. **B**. Analogous to panel A, but using the optimal AWSEM-IF blend models.

**Figure S5.**
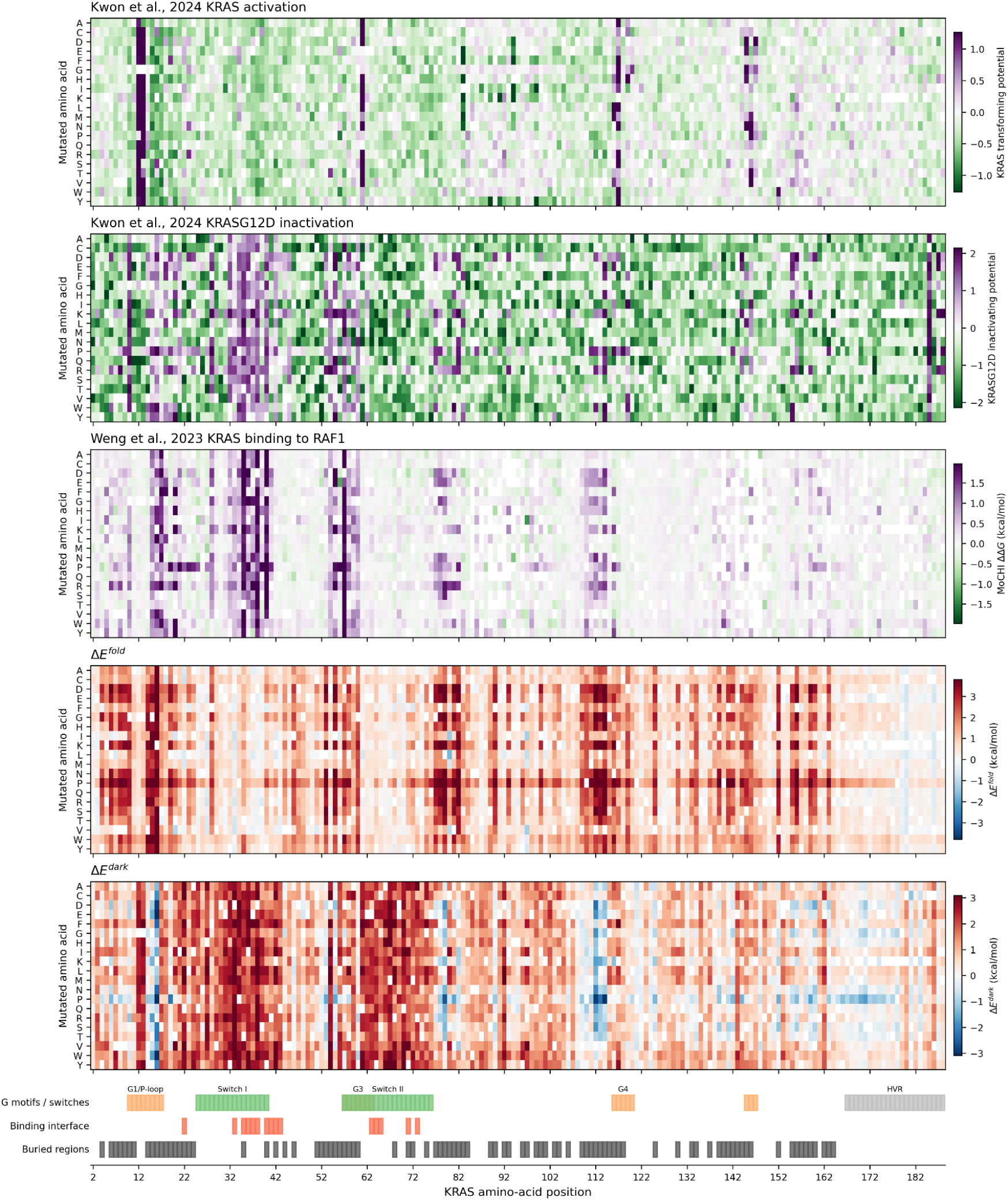
KRAS experimental Deep Mutational Scannings and model predictions for folding-stability and dark energy changes. The top panels show Kwon et al. [49] experimental assays for KRAS wildtype oncogenic activation and for the KRAS common cancer mutant G12D inactivation, respectively. The middle panel shows Weng et al. [8] results for KRAS binding energy to RAF1. The bottom panels show the folding and dark energy change predictions. Functional and structural annotations are shown at the bottom. Buried regions were defined as residues with relative solvent-accessible surface area (RSA) < 0.25 on the corresponding AlphaFold structure.

**Figure S6.**
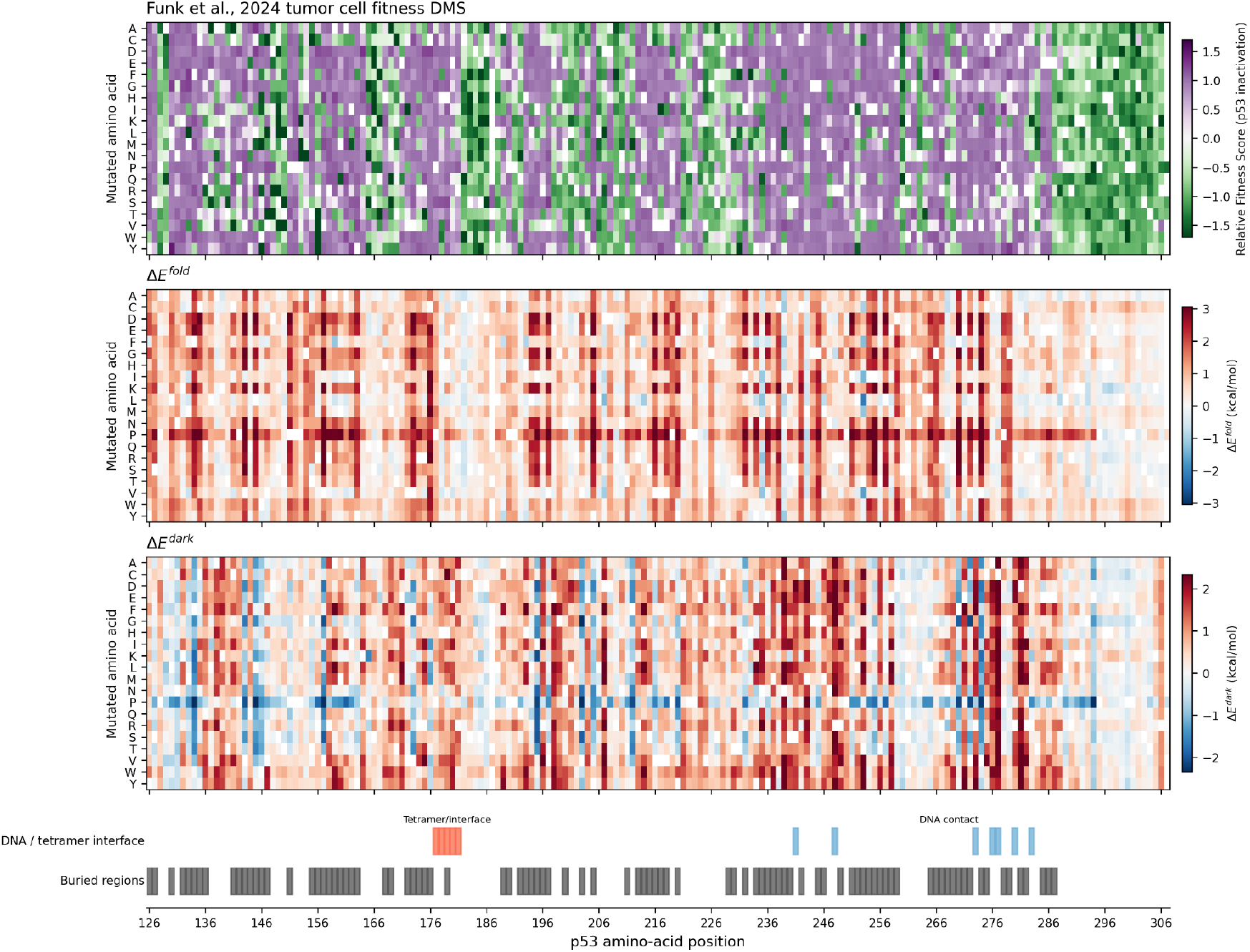
p53 experimental Deep Mutational Scanning and model predictions for folding-stability and dark energy changes. The top panel shows Funk et al. [50] tumor cell fitness experimental measurement for p53. The middle and bottom panels show the folding and dark energy change predictions. Functional and structural annotations are shown at the bottom. Buried regions were defined as residues with relative solvent-accessible surface area (RSA) < 0.25 on the corresponding AlphaFold structure.

## References

1. Bryngelson, J. D. & Wolynes, P. G. Spin glasses and the statistical mechanics of protein folding. Proc. Natl. Acad. Sci. 84, 7524–7528 (1987).

2. Ramanathan, S. & Shakhnovich, E. Statistical mechanics of proteins with ‘“evolutionary selected”’ sequences. Phys. Rev. E 50, 1303–1312 (1994).

3. Ferreiro, D. U., Komives, E. A. & Wolynes, P. G. Frustration in biomolecules. Q. Rev. Biophys. 47, 285–363 (2014).

4. Beadle, B. M. & Shoichet, B. K. Structural Bases of Stability–function Tradeoffs in Enzymes. J. Mol. Biol. 321, 285–296 (2002).

5. Notin, P. et al. ProteinGym: Large-Scale Benchmarks for Protein Fitness Prediction and Design. in Advances in Neural Information Processing Systems vol. 36 64331–64379 (2023).

6. Sánchez, I. E., Tejero, J., Gómez-Moreno, C., Medina, M. & Serrano, L. Point Mutations in Protein Globular Domains: Contributions from Function, Stability and Misfolding. J. Mol. Biol. 363, 422–432 (2006).

7. Gerasimavicius, L., Livesey, B. J. & Marsh, J. A. Loss-of-function, gain-of-function and dominant-negative mutations have profoundly different effects on protein structure. Nat. Commun. 13, 3895 (2022).

8. Weng, C., Faure, A. J., Escobedo, A. & Lehner, B. The energetic and allosteric landscape for KRAS inhibition. Nature 626, 643–652 (2024).

9. Faure, A. J. et al. Mapping the energetic and allosteric landscapes of protein binding domains. Nature 604, 175–183 (2022).

10. Beltran, A., Naqvi, M. M., Faure, A. J. & Lehner, B. The allosteric landscape of the Src kinase. Sci. Adv. 12, eaea2726 (2026).

11. Coyote-Maestas, W., Nedrud, D., He, Y. & Schmidt, D. Determinants of trafficking, conduction, and disease within a K+ channel revealed through multiparametric deep mutational scanning. eLife 11, e76903 (2022).

12. Martí-Aranda, A. & Lehner, B. Seven complete comparative maps of allosteric mutations in a protein family. Nat. Commun. 17, 5487 (2026).

13. Vanella, R. et al. Understanding activity-stability tradeoffs in biocatalysts by enzyme proximity sequencing. Nat. Commun. 15, 1807 (2024).

14. Galpern, E. A., Bueno, C., Sánchez, I. E., Wolynes, P. G. & Ferreiro, D. U. Probing the dark energy in the functional protein universe. Proc. Natl. Acad. Sci. 123, e2531111123 (2026).

15. Cagiada, M., Jonsson, N. & Lindorff-Larsen, K. Decoding molecular mechanisms for loss-of-function variants in the human proteome. Preprint at 10.7554/eLife.108160.1 (2025).

16. Liao, X. & Lehner, B. Allostery is a widespread cause of loss-of-function variant pathogenicity. Nat. Commun. (2026).

17. Hsu, C. et al. Learning inverse folding from millions of predicted structures. in Proceedings of the 39th International Conference on Machine Learning vol. 162 8946–8970 (2022).

18. Dauparas, J. et al. Robust deep learning–based protein sequence design using ProteinMPNN. Science 378, 49–56 (2022).

19. Frellsen, J. et al. Zero-shot protein stability prediction by inverse folding models: a free energy interpretation. in Advances in Neural Information Processing Systems (eds Belgrave, D.et al.) vol. 38 84641–84667 (Curran Associates, Inc., 2025).

20. Dutton, O. et al. Improving Inverse Folding models at Protein Stability Prediction without additional Training or Data. Preprint at 10.1101/2024.06.15.599145 (2024).

21. Birnbaum, F. & Keating, A. E. Beyond native sequence recovery: Improved modeling of the sequence-energy landscape of protein structures. Proc. Natl. Acad. Sci. 123, e2535494123 (2026).

22. Shuai, R. W., Lu, T., Bhatti, S., Kouba, P. & Huang, P.-S. Ensemble-conditioned protein sequence design with Caliby. Preprint at 10.1101/2025.09.30.679633 (2025).

23. Ferreiro, D. U., Hegler, J. A., Komives, E. A. & Wolynes, P. G. Localizing frustration in native proteins and protein assemblies. Proc. Natl. Acad. Sci. 104, 19819–19824 (2007).

24. Freiberger, M. I. et al. Local energetic frustration conservation in protein families and superfamilies. Nat. Commun. 14, 8379 (2023).

25. Listgarten, J. & Jiang, H. How artificial intelligence is reengineering protein engineering. Science 392, 159–166 (2026).

26. Davtyan, A. et al. AWSEM-MD: Protein structure prediction using coarse-grained physical potentials and bioinformatically based local structure biasing. J. Phys. Chem. B 116, 8494–8503 (2012).

27. Freiberger, M. I., Guzovsky, A. B., Wolynes, P. G., Parra, R. G. & Ferreiro, D. U. Local frustration around enzyme active sites. Proc. Natl. Acad. Sci. 116, 4037–4043 (2019).

28. Galpern, E. A., Jaafari, H., Bueno, C., Wolynes, P. G. & Ferreiro, D. U. Reassessing the exon–foldon correspondence using frustration analysis. Proc. Natl. Acad. Sci. 121, e2400151121 (2024).

29. Tokuriki, N., Stricher, F., Schymkowitz, J., Serrano, L. & Tawfik, D. S. The Stability Effects of Protein Mutations Appear to be Universally Distributed. J. Mol. Biol. 369, 1318–1332 (2007).

30. Miyazawa, S. Selection originating from protein stability/foldability: Relationships between protein folding free energy, sequence ensemble, and fitness. J. Theor. Biol. 433, 21–38 (2017).

31. Tsuboyama, K. et al. Mega-scale experimental analysis of protein folding stability in biology and design. Nature 620, 434–444 (2023).

32. Meier, J. et al. Language models enable zero-shot prediction of the effects of mutations on protein function. in Advances in Neural Information Processing Systems vol. 34 29287–29303 (2021).

33. Morcos, F., Schafer, N. P., Cheng, R. R., Onuchic, J. N. & Wolynes, P. G. Coevolutionary information, protein folding landscapes, and the thermodynamics of natural selection. Proc. Natl. Acad. Sci. 111, 12408–12413 (2014).

34. Dieckhaus, H., Brocidiacono, M., Randolph, N. Z. & Kuhlman, B. Transfer learning to leverage larger datasets for improved prediction of protein stability changes. Proc. Natl. Acad. Sci. 121, e2314853121 (2024).

35. Beltran, A., Jiang, X., Shen, Y. & Lehner, B. Site-saturation mutagenesis of 500 human protein domains. Nature 637, 441–449 (2025).

36. Furnham, N. et al. The Catalytic Site Atlas 2.0: cataloging catalytic sites and residues identified in enzymes. Nucleic Acids Res. 42, D485–D489 (2014).

37. Stojanoski, V. et al. Removal of the Side Chain at the Active-Site Serine by a Glycine Substitution Increases the Stability of a Wide Range of Serine β-Lactamases by Relieving Steric Strain. Biochemistry 55, 2479–2490 (2016).

38. Thomas, V. L., McReynolds, A. C. & Shoichet, B. K. Structural Bases for Stability–Function Tradeoffs in Antibiotic Resistance. J. Mol. Biol. 396, 47–59 (2010).

39. Marchler-Bauer, A. et al. CDD: NCBI’s conserved domain database. Nucleic Acids Res. 43, D222–D226 (2015).

40. Martínez-Jiménez, F. et al. A compendium of mutational cancer driver genes. Nat. Rev. Cancer 20, 555–572 (2020).

41. Bishop, J. M. Molecular themes in oncogenesis. Cell 64, 235–248 (1991).

42. Hanahan, D. & Weinberg, R. A. The Hallmarks of Cancer. Cell 100, 57–70 (2000).

43. Vogelstein, B. & Kinzler, K. W. Cancer genes and the pathways they control. Nat. Med. 10, 789–799 (2004).

44. Zhao, M., Kim, P., Mitra, R., Zhao, J. & Zhao, Z. TSGene 2.0: an updated literature-based knowledgebase for tumor suppressor genes. Nucleic Acids Res. 44, D1023–D1031 (2016).

45. Bleeker, F. E. et al. The prognostic IDH1 R132 mutation is associated with reduced NADP+-dependent IDH activity in glioblastoma. Acta Neuropathol. (Berl.) 119, 487–494 (2010).

46. Kandoth, C. et al. Mutational landscape and significance across 12 major cancer types. Nature 502, 333–339 (2013).

47. Prior, I. A., Lewis, P. D. & Mattos, C. A Comprehensive Survey of Ras Mutations in Cancer. Cancer Res. 72, 2457–2467 (2012).

48. Joerger, A. C. & Fersht, A. R. Structure–function–rescue: the diverse nature of common p53 cancer mutants. Oncogene 26, 2226–2242 (2007).

49. Kwon, J. J. et al. Comprehensive structure-function analysis reveals gain- and loss-of-function mechanisms impacting oncogenic KRAS activity. Preprint at 10.1101/2024.10.22.618529 (2024).

50. Funk, J. S. et al. Deep CRISPR mutagenesis characterizes the functional diversity of TP53 mutations. Nat. Genet. 57, 140–153 (2025).

51. Sevim Bayrak, C. et al. Identification of discriminative gene-level and protein-level features associated with pathogenic gain-of-function and loss-of-function variants. Am. J. Hum. Genet. 108, 2301–2318 (2021).

52. Yan, D. & Ishihara, K. Two Kir2.1 channel populations with different sensitivities to Mg2+ and polyamine block: a model for the cardiac strong inward rectifier K+ channel. J. Physiol. 563, 725–744 (2005).

53. Lu, C.-W. et al. Functional and clinical characterization of a mutation in KCNJ2 associated with Andersen-Tawil syndrome. J. Med. Genet. 43, 653–659 (2006).

54. Plaster, N. M. et al. Mutations in Kir2.1 Cause the Developmental and Episodic Electrical Phenotypes of Andersen’s Syndrome. Cell 105, 511–519 (2001).

55. Tristani-Firouzi, M. et al. Functional and clinical characterization of KCNJ2 mutations associated with LQT7 (Andersen syndrome). J. Clin. Invest. 110, 381–388 (2002).

56. Hattori, T. et al. A novel gain-of-function KCNJ2 mutation associated with short-QT syndrome impairs inward rectification of Kir2.1 currents. Cardiovasc. Res. 93, 666–673 (2012).

57. Deo, M. et al. KCNJ2 mutation in short QT syndrome 3 results in atrial fibrillation and ventricular proarrhythmia. Proc. Natl. Acad. Sci. 110, 4291–4296 (2013).

58. Orenbuch, R. et al. Proteome-wide model for human disease genetics. Nat. Genet. 57, 3165–3174 (2025).

59. Cheng, J. et al. Accurate proteome-wide missense variant effect prediction with AlphaMissense. Science 381, eadg7492 (2023).

60. Gao, H. et al. The landscape of tolerated genetic variation in humans and primates. Science 380, eabn8153 (2023).

61. Song, H. et al. Diverse rescue potencies of p53 mutations to ATO are predetermined by intrinsic mutational properties. Sci. Transl. Med. 15, eabn9155 (2023).

62. Stein, A., Fowler, D. M., Hartmann-Petersen, R. & Lindorff-Larsen, K. Biophysical and Mechanistic Models for Disease-Causing Protein Variants. Trends Biochem. Sci. 44, 575–588 (2019).

63. Mighell, T. L. & Lehner, B. A small molecule stabilizer rescues the surface expression of nearly all missense variants in a GPCR. Nat. Struct. Mol. Biol. 32, 2429–2440 (2025).

64. Schafer, N. P., Kim, B. L., Zheng, W. & Wolynes, P. G. Learning To Fold Proteins Using Energy Landscape Theory. Isr. J. Chem. 54, 1311–1337 (2014).

65. Hou, Q., Rooman, M. & Pucci, F. Enzyme Stability-Activity Trade-Off: New Insights from Protein Stability Weaknesses and Evolutionary Conservation. J. Chem. Theory Comput. 19, 3664–3671 (2023).

66. Figliuzzi, M., Jacquier, H., Schug, A., Tenaillon, O. & Weigt, M. Coevolutionary landscape inference and the context-dependence of mutations in beta-lactamase TEM-1. Mol. Biol. Evol. 33, 268–280 (2016).

67. Poley-Gil, M. et al. Adaptive and Spandrel-like Constraints at Functional Sites in Protein Folds. Preprint at 10.64898/2026.02.09.704872 (2026).

68. Weinstein, E., Amin, A., Frazer, J. & Marks, D. Non-identifiability and the Blessings of Misspecification in Models of Molecular Fitness. in Advances in Neural Information Processing Systems vol. 35 5484–5497 (2022).

69. Pugh, C. W. J., Nuñez-Valencia, P. G., Dias, M. & Frazer, J. From Likelihood to Fitness: Improving Variant Effect Prediction in Protein and Genome Language Models. in Advances in Neural Information Processing Systems vol. 38 130835–130866 (2025).

70. Gordon, C., Lu, A. X. & Abbeel, P. Protein Language Model Fitness Is a Matter of Preference. in The Thirteenth International Conference on Learning Representations (2024).

71. Chillón-Pino, D., Badonyi, M., Semple, C. A. & Marsh, J. A. Protein structural context of cancer mutations reveals molecular mechanisms and candidate driver genes. Cell Rep. 43, 114905 (2024).

72. Stehr, H. et al. The structural impact of cancer-associated missense mutations in oncogenes and tumor suppressors. Mol. Cancer 10, 54 (2011).

73. Batyuk, A., Wu, Y., Honegger, A., Heberling, M. M. & Plückthun, A. DARPin-based crystallization chaperones exploit molecular geometry as a screening dimension in protein crystallography. J. Mol. Biol. 428, 1574–1588 (2016).

74. Cianferoni, D. et al. Artificial intelligence and first-principle methods in protein redesign: A marriage of convenience? Protein Sci. 34, e70210 (2025).

75. Varadi, M. et al. AlphaFold Protein Structure Database: massively expanding the structural coverage of protein-sequence space with high-accuracy models. Nucleic Acids Res. 50, D439–D444 (2022).

76. Jumper, J. et al. Highly accurate protein structure prediction with AlphaFold. Nature 596, 583–589 (2021).

77. Muiños, F., Martínez-Jiménez, F., Pich, O., Gonzalez-Perez, A. & Lopez-Bigas, N. In silico saturation mutagenesis of cancer genes. Nature 596, 428–432 (2021).

78. Martincorena, I. et al. Universal Patterns of Selection in Cancer and Somatic Tissues. Cell 171, 1029–1041.e21 (2017).

79. Bueno, C., Jaafari, H., Galpern, E. A., Ferreiro, D. U. & Wolynes, P. G. Python Frustratometer. https://github.com/HanaJaafari/Frustratometer/tree/galpern2025 (2026).

